# Matrix topology guides collective cell migration *in vivo*

**DOI:** 10.1101/2022.01.31.478442

**Authors:** Karen G. Soans, Ana Patricia Ramos, Jaydeep Sidhaye, Abhijeet Krishna, Anastasia Solomatina, Karl B. Hoffmann, Raimund Schlüßler, Jochen Guck, Ivo F. Sbalzarini, Carl D. Modes, Caren Norden

## Abstract

Diverse modes of cell migration shape organisms in health and disease and much research has focused on the role of intracellular and extracellular components in different cell migration phenomena. What is less explored, however, is how the arrangement of the underlying extracellular matrix that many cells move upon *in vivo* influences migration.

Combining novel transgenic lines and image analysis pipelines, reveals that during zebrafish optic cup formation cells use cryptopodia-like protrusions to migrate collectively and actively over a topologically changing matrix. These changing topologies correspond to different cell-matrix interactions. Interference with matrix topology results in loss of cryptopodia and inefficient migration. Thus, matrix topology influences the efficiency of directed collective cell migration during eye morphogenesis, a concept likely conserved in other developmental and disease contexts.

**One-Sentence Summary:** Dynamic cell-matrix interactions, crucial for successful collective rim cell migration, rely on extracellular matrix topologies during optic cup development *in vivo*.

## Main Text

During organogenesis and in homeostasis, cells often actively move to the location where they later function (*1, 2*). Further, many cancers are worsened when cell migration causes metastasis (*3*).

So far, most studies have focused on understanding cell migration during development, as findings are often translatable to disease states (*4*). Cell migration during development occurs via distinct modes broadly classified as single or collective cell migration (*5*). Singly migrating cells show the intrinsic ability to move in a certain direction as shown for primordial germ cell migration (*1*) or neuronal migration (*6*). Collectively migrating cells move as a cohesive group in which they maintain contact with each other to coordinate their movements while they are additionally steered by environmental factors (*4, 7*). Examples of collective cell migration include follicle epithelial cells that shape the *Drosophila* egg chamber (*8*) or the migrating lateral line primordium in zebrafish (*9*).

All types of cell migration depend on both intrinsic and extrinsic factors for directionality and migration efficiency. The role of cell intrinsic factors like adherens junctions, cytoskeletal elements, and basal focal adhesions (FA) have been studied in depth and shown to be essential to facilitate cell migration in numerous contexts (*10, 11*). Further, cell extrinsic factors like growth factors or guiding chemokines provide directionality for long-range single and collective cell migration (*1, 9*).

In addition to cell intrinsic factors and extrinsic chemical signaling, it is now clear that physical parameters like tissue stiffness, substrate density, or topology (*12, 13*) can also be instructive for cell migration. However, here, many details are still unknown. In particular, the role of the extracellular matrix (ECM), that is encountered by many migrating cells *in vivo* is still underexplored (*14*). To understand how the ECM could influence collective cell migration, *in vitro* studies in which the density, stiffness and arrangement patterns of the ECM mimicked the *in vivo* situation have revealed a spectrum of possibilities by which ECM properties can influence cell migration. This has included ECM stiffness (*15, 16*) and topology, which involves the porosity, alignment and density of the matrix (*17*). So far, however, the extent to which ECM properties affect cell migration *in vivo* is not yet fully explored. One reason is that here the dynamic visualization of cell-matrix interactions has been challenging due to the lack of appropriate tools.

The developing optic cup in zebrafish is a model that allows for such visualization as it is situated at the outside of the developing embryo and is approachable to quantitative long-term imaging. During optic cup formation (OCF), collectively migrating rim cells move within a continuous epithelium to shape the hemispherical tissue (*18, 19*). Rim cells require an intact ECM and cell-matrix interactions for successful migration (*18*) but the exact interplay between ECM and rim cells is unknown. We investigate this phenomenon by introducing tools to visualize the ECM *in vivo*, combined with new analysis pipelines to quantify cell-matrix interactions. We find that rim cells migrate collectively over a uniformly compressible ECM. The topology of the matrix varies along the migration path and influences cell-matrix interactions and migration dynamics. These interactions rely on basal cryptopodia-like protrusions, basal protrusions that extend beneath the next cell along the direction of migration. When the topology of the matrix is perturbed, these cryptopodia are lost and migrating rim cells become less directed. Thus, we directly link matrix topology to the guidance of collective cell migration *in vivo* generating an important reference to investigate similar phenomena in other developmental and disease contexts.

### Laminin does not undergo diffusive turnover in the optic cup ECM

To reveal possible interactions between collectively migrating rim cells and the underlying matrix, we generated markers to visualize the ECM during OCF *in vivo*. As laminin knockdown impairs efficient rim migration (*18*), transgenic laminin lines were generated using a bacterial artificial chromosome (BAC) containing the laminin α-1 gene along with its native gene regulatory elements (lama1 in zebrafish) and a large upstream region to preserve possible regulatory elements (for details see Materials and Methods). This ensured close-to-endogenous expression of laminin. Laminin was tagged with the fluorescent proteins sf-GFP, mKate2 and photoconvertible dendra (Fig. 1A and fig. S1B). Expression of these constructs matched laminin α-1 antibody staining (Fig. 1B). Further, transplantation of *Tg(lama1:lama1-sfGFP)* cells into *Tg(lama1:lama1-mKate2)* showed that mKate2 and GFP expression colocalize (fig. S1C). Transplantations also revealed that the laminin deposited around the optic cup arises from more than one tissue as even when transplanted cells only appeared outside of the optic cup, they contributed to laminin in the optic cup ECM (fig. S1C and Table S6, N = 3/4 embryos).

**Fig. 1.**
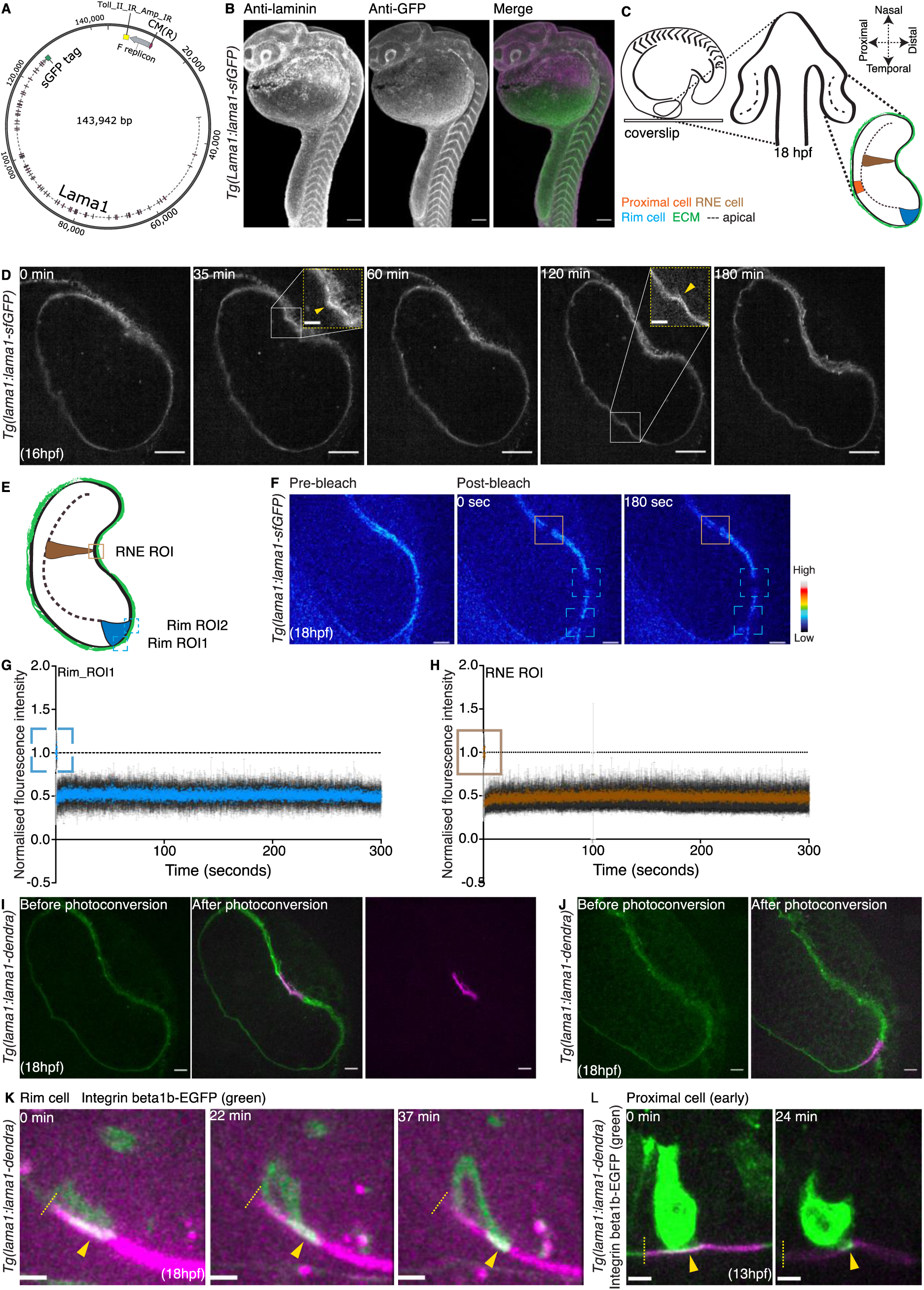
Laminin does not diffuse freely within the matrix and rim cells actively migrate over the ECM. (**A**) Schematic of zebrafish *lama1* BAC (CH211-170M3) tagged with the fluorescent protein super-folded GFP (sfGFP). (**B**) Snapshots of fixed *Tg(lama1:lama1-sfGFP)* zebrafish embryos stained against GFP (green) and laminin (laminin α-1) (gray) antibodies at 24hpf (scale bar = 100 μm). (**C**) Schematic depicting the mounting and imaging strategy for zebrafish embryos at 18hpf. (**D**) Montage of live *Tg(lama1:lama1-sfGFP)* embryos mounted at 16hpf (scale bar = 30 μm). Insets indicate deformations of laminin (yellow arrows) during optic cup formation (OCF) (scale bar = 10 μm). (**E**) Schematic of the zebrafish optic cup at 18hpf indicating the region of interest (ROI) in which bleaching was performed. (**F**) Montage of live *Tg(lama1:lama1-sfGFP)* embryos (royal LUT) at 18hpf pre and post bleaching (scale bar = 10 μm). (**G-H**) Normalized fluorescence intensity plots of laminin in the bleached ROIs of rim and RNE (mean and SD, n = 7 embryos per experiment). (**I-J**) Snapshots of live *Tg(lama1:lama1-dendra)* embryos at 18hpf in which laminin tagged to dendra was photoconverted from green to magenta in the RNE and rim regions (scale bar = 10 μm). (**K-L**) Montage of focal adhesion (FA) dynamics in live *Tg(lama1:lama1-dendra)* embryos at 18hpf and 13hpf (scale bar = 5 μm). FAs of rim cells are labeled with Integrin beta1b-EGFP (green) and photoconverted laminin is seen in magenta. Dashed yellow lines indicate the start of the photoconverted laminin region. Yellow arrowheads point to moving FAs.

Time lapse imaging of *Tg(lama1:lama1-sfGFP)* between 16hpf and 19hpf, the stages at which OCF occurs, revealed that the ECM surrounding the optic cup undergoes frequent deformations (Fig. 1D and movie S1) indicating some matrix dynamicity. To understand whether these deformations are accompanied by matrix turnover, fluorescence recovery after photobleaching (FRAP) was performed. Over short time spans (5min imaged at 50ms intervals) the intensity of laminin did not recover (Fig. 1, E to H, fig. S1D and movie S2). Over longer time periods in the range of hours however, a gradual recovery of laminin was observed, most likely due to the deposition of new protein into the matrix (fig. S1, E to G and movie S3).

Thus, laminin does not freely diffuse in the optic cup ECM over short time scales and can be deposited by cells outside the optic cup.

### Rim cells actively migrate over the underlying ECM using cryptopodia like protrusions

The low laminin turnover rate suggested that this structure served as a stable scaffold for rim cells to actively move over. Another possibility was that the whole matrix was moving towards the retinal neuroepithelium (RNE) thereby passively dragging rim cells with it, as seen previously in other contexts (*20*). To differentiate between these two possibilities, *Tg(lama1:lama1-dendra)* embryos (Fig. 1I and fig. S1H) were photoconverted in the rim region (Fig. 1, J to K) and focal adhesions (FAs) were labeled by Integrin beta1b-EGFP. This allowed the tracking of rim cells relative to matrix movements (movie S4) and showed that rim cells moved actively over the matrix (Fig. 1K). The same experiment was conducted at earlier stages of OCF (13hpf – 14hpf), revealing that even before obvious RNE invagination onset, rim and proximal cells showed active movement over the matrix towards the prospective RNE (Fig. 1L, fig. S1I and movie S4).

As it was shown that rim cell migration correlated with protrusive activity (*18*), we investigated the ultrastructure of the protrusions involved through transmission electron microscopy (TEM). We found that protrusions did not resemble filopodia or lamellipodia as seen for other collectively migrating cells (*11, 21*) but instead showed cryptopodia characteristics as they extended beneath the cell in front, along the direction of migration (*22*). Such cryptopodia have previously been shown to direct collective cell migration *in vitro*, in systems that lack a leading edge or leader cells (*23*) similar to the optic cup (*7*). Interestingly, the cryptopodia-like protrusions are less extended in cells already positioned close to the distal side of the tissue. Further, cells in the RNE no longer extended cryptopodia in any particular direction (Fig. 2A to C and fig. S2E).

**Fig. 2.**
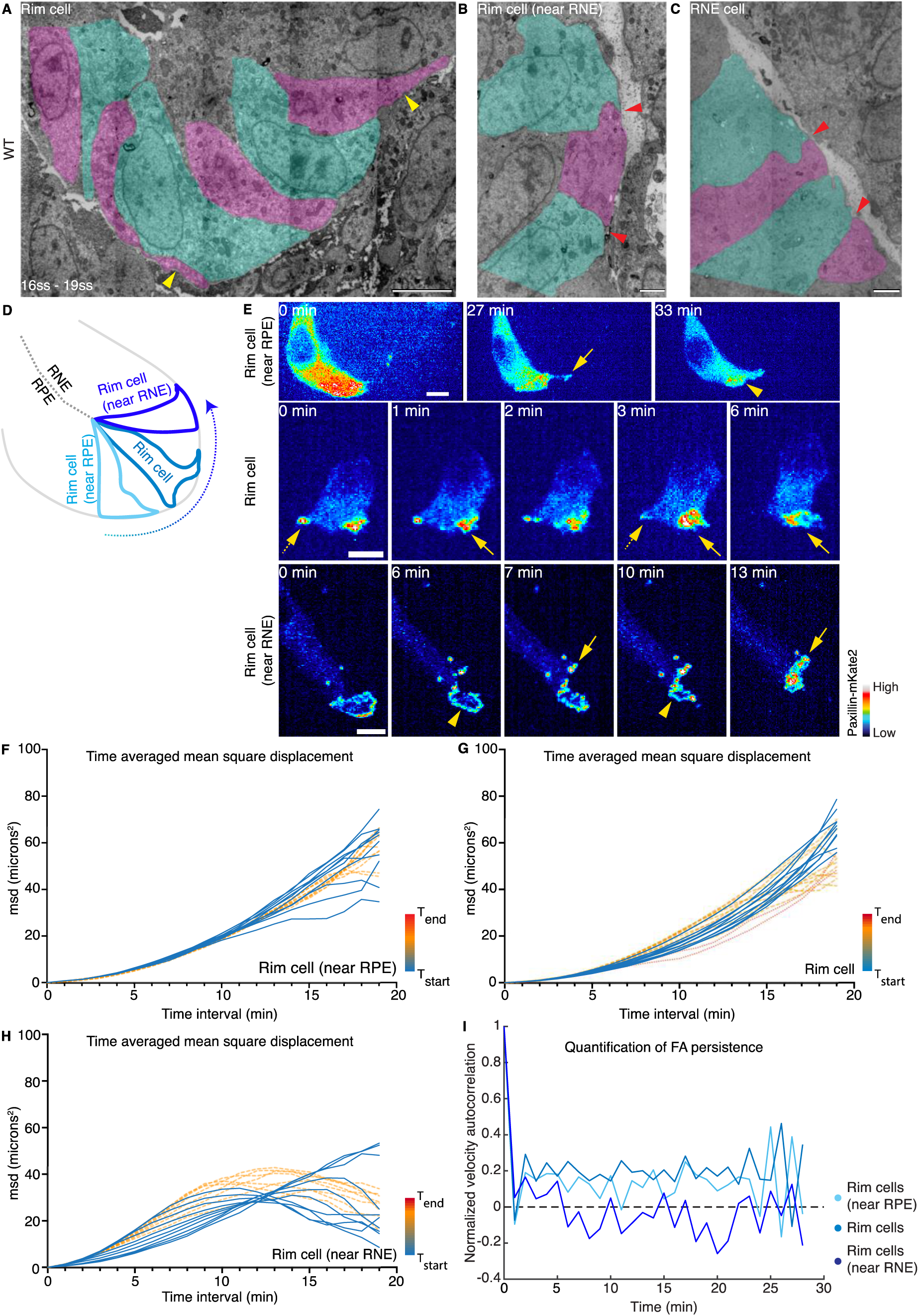
Rim cells exhibit cryptopodia like protrusions and different FA dynamics depending on the site of migration. (**A-C**) TEM images of basal protrusions of migrating cells in WT embryos covering the rim region (scale bar = 5 μm), near the RNE (scale bar = 2 μm) and in the RNE region (scale bar = 2 μm) between 16ss and 19ss. Yellow arrowheads indicate cryptopodia like protrusions. Red arrows indicate smaller, less extended, protrusions in the rim cells near the RNE and in the RNE region. Green and pink areas delineate single cells. (**D**) Schematic of optic cup at 18hpf depicting migrating rim cells near the RPE, RNE and in the center of the rim region. (**E**) Montage of migrating cells in the different regions of the rim. focal adhesions are labeled with Paxillin-mKate2 (royal LUT). Yellow arrows point to newly forming leading FA, yellow arrowheads point to sliding FA. Dotted yellow arrow points to detaching FA (scale bar = 5 μm). (**F-H**) Time averaged mean square displacement (tamsd) plots of sub tracks of migrating rim cells near RPE, in the rim region and near the RNE (means plotted). Color code from blue to red indicates increasing time from the start to the end of the movie. (**I**) Normalized velocity autocorrelation plots (means plotted) of migrating rim cells near the RPE (n = 7 cells), RNE (n = 3 cells) and in the center of the rim region (n = 7 cells).

Thus, active rim cell movement starts at very early stages of OCF, before the onset of tissue invagination and rim migration and is accompanied by cryptopodia formation with different morphologies depending on rim cell location.

### Rim cells exhibit different focal adhesion dynamics in different regions of the migratory path

We speculated that the different morphologies of basal cryptopodia along the path of migration were linked to different cell interactions with the underlying ECM and tested this idea via live imaging. Mosaic labeling of the FA proteins paxillin (Paxillin-mKate2) or integrin (Integrin beta1b-EGFP, Integrin beta1b-mKate2) in combination with the transgenic laminin lines revealed that FA behavior changed from dynamic to more stable once rim cells reached the inner side of the neuroepithelium (movie S5). In particular, FA interactions with the underlying matrix changed along the migration path and could be divided into three categories (Fig. 2, D to E and fig. S2A). **Category 1:** The majority of migrating cells in the rim region showed FAs that detached from and reattached to laminin (Fig. 2E dotted yellow arrow) and thus rim cells moved in a ‘step-like’ manner (fig. S2D). **Category 2:** FAs of migrating cells near the RPE and migrating cells near the RNE (Fig. 2D) interacted more continuously with laminin (Fig. 2E yellow arrow) and moved in a ‘sliding’ manner towards the RNE (fig. S2D). **Category 3**: For cells in the RNE, most of the observed FAs interacted continuously with the underlying laminin, ‘wiggling’ back and forth without considerably changing their position (fig. S2, B to D). (For additional visualization of these phenomena refer to kymographs in fig. 2A and supplement movie S5).

To analyze FA dynamics in more quantitative detail, a segmentation and tracking tool was set up (*24*) that allowed the segmentation of only the FA signal that colocalized with the underlying laminin matrix (see Materials and Methods) (movie S6). This tool enabled the calculation of the time averaged mean square displacement (tamsd) and velocity autocorrelation, allowing for the analysis of directionality and persistence of FA movements. The positive curvature of the tamsd graphs showed that FAs of migrating cells close to the RPE and in the rim region underwent directed movements (Fig. 2, F to G and fig. S2, F to G). Interestingly, FAs of migrating cells close to the RNE displayed less directed movements and migrated in the least persistent manner (Fig. 2, H to I and fig. S2H). This is in line with the qualitative analysis that showed that these cells step back more often than migrating cells in the rim region and near the RPE (fig. S2D).

Overall, qualitative and quantitative evaluation of FA dynamics over the matrix showed that rim cells transition from directed to less directed movement prior to stopping when entering the RNE (fig. S2, B to C and fig. S2I).

### Changing focal adhesion dynamics are accompanied by changes in laminin topology

*In vitro* studies showed that changes of ECM properties like stiffness, density and arrangement of fibrils can result in changing FA dynamics (*17, 25, 26*). Thus, we characterized and quantified matrix properties in the different migratory zones.

To assess matrix compressibility, Brillouin microscopy, a non-invasive method that measures the compressibility of tissues and matrices *in vivo*, was used (*27*). Measurements were carried out in the rim and RNE region of the developing optic cup at 18hpf (fig. S3, A to E). Statistical analysis of the data revealed no significant difference in compressibility between the two regions (see Materials and Methods, fig. S3, A to E). More so, even in the few embryos for which differences between rim-ECM and RNE-ECM matrix were detected, no common trend emerged (fig. S3, D to E). Thus, no consistent matrix compressibility gradient exists making matrix stiffness an unlikely factor to drive changing interactions between rim cells and the ECM.

High resolution confocal imaging of laminin α-1 staining revealed that laminin arrangements differed between the rim and RNE regions. While laminin in the RNE region appeared as aligned fibrils, in the rim region laminin was more porous and the fibrils less aligned (Fig. 3A).

**Fig. 3.**
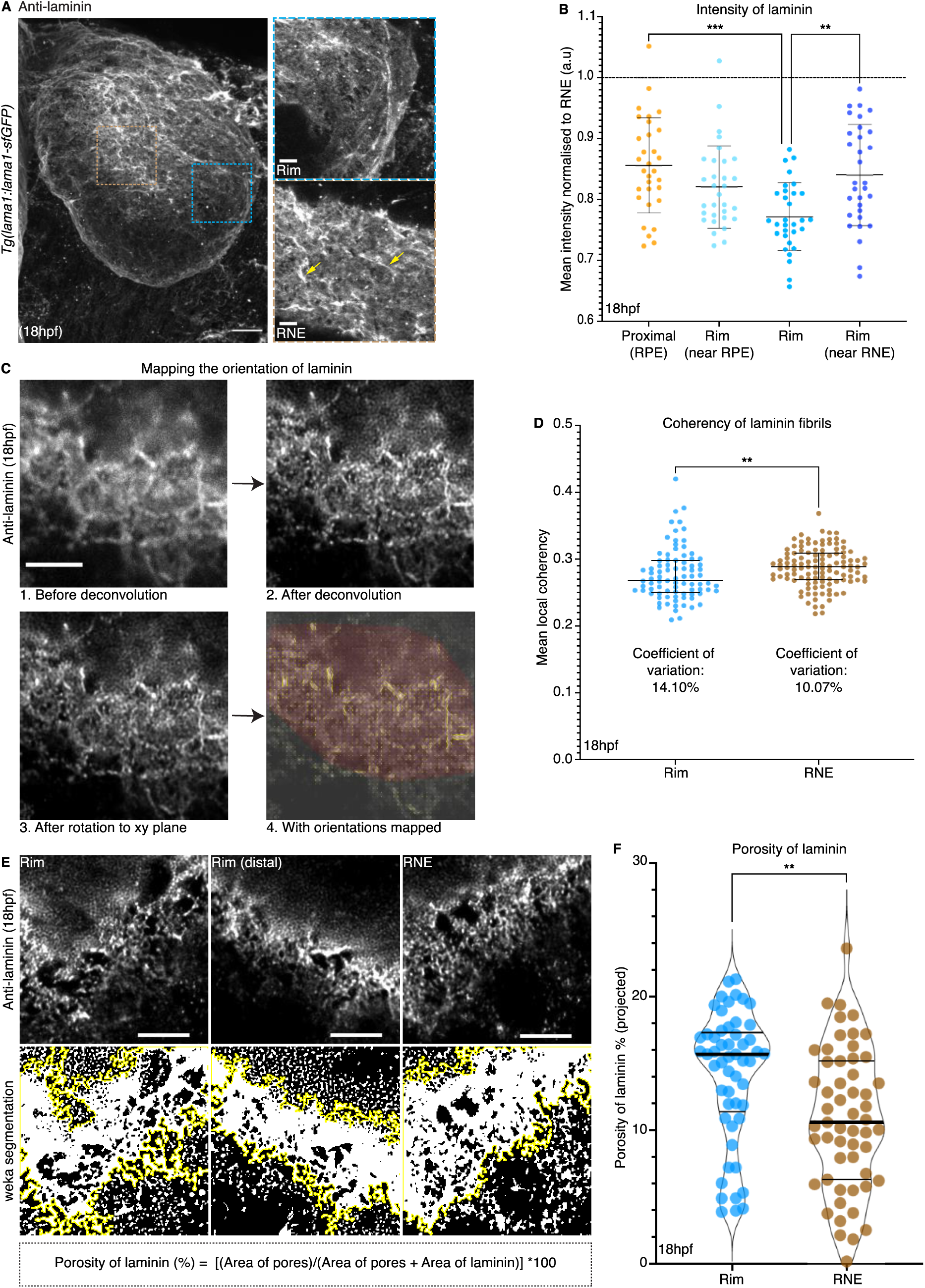
Laminin shows different topologies in rim compared to RNE regions. (**A**) Maximum intensity projection of *Tg(lama1:lama1-sfGFP)* embryos at 18hpf stained against laminin α-1 (gray, scale bar = 20 μm). Brown and blue dotted boxes mark the region of RNE and rim in the inlay. Yellow arrows point to fibrils of laminin (scale bar = 5 μm). (**B**) Laminin mean intensity graph of the proximal and rim regions normalized to the mean intensity of laminin in the RNE region (n = 6 embryos). Kruskal-Wallis test with Dunn’s multiple comparison test (mean with SD, ** p = 0.0022, *** p = 0.0001). (**C**) Image analysis pipeline to quantify the mean local orientation of laminin fibrils. Maximum intensity projections of *Tg(Vsx2:GFP)* embryos stained against laminin α-1 (scale bar = 5 μm). (**D**) Graph of mean local coherency (defined in Materials and Methods) of laminin fibrils in the rim (n = 7 embryos) and RNE region (n = 8 embryos). Two tailed Mann Whitney test (median with interquartile range, ** p = 0.0021). (**E**) Maximum intensity projections of ROIs used to measure the porosity of laminin including the formula used to calculate the porosity of laminin (scale bar = 5 μm). (**F**) Graph of porosity of laminin (% projected) in the rim (n = 8 embryos) and RNE (n = 8 embryos) region. Kruskal-Wallis test with Dunn’s multiple comparison test (median with quartiles, ** p = 0.0010).

We thus asked whether overall matrix topology could influence rim migration as changes in substrate topology and density have the ability to influence FA dynamics *in vitro* (*17, 25, 26*). To investigate the topology of the matrix, we established image analysis pipelines (for details refer to Materials and Methods) that quantified the arrangement, porosity and intensity of laminin in different optic cup regions. This analysis showed that the orientation of laminin fibrils was more coherent in the RNE region compared to the rim region (Fig. 3, C to D) at 18hpf and became progressively more aligned in the RNE region at 19hpf (fig. S3, I to J). In addition, the porosity of laminin was much higher in the rim region compared to the RNE at 18hpf (Fig. 3, E to F for analysis details refer to Materials and Methods). Interestingly, the degree of porosity within the rim region decreased closer to the RNE (fig. S3H). Intensity measurements of laminin using the *Tg(lama1:lama1-sfGFP)* and *Tg(lama1::lama-mKate2)* lines concurred with the porosity measurements as laminin intensity was the highest in the RNE and lowest in the rim region at 18hpf (Fig. 3B and fig. S3F).

Overall, this showed that the topology of laminin is spatially distinct in different regions of the optic cup suggesting a possible link between matrix topology and the different FA dynamics observed.

### Differences in topology result in changing physical properties of laminin

Changes in matrix arrangement can lead to changing mechanical properties of the matrix, which in turn influence cell-matrix interactions and cell migration as shown *in vitro* and *in vivo* (*28, 26*). While we did not see significant changes in overall compressibility between the rim- ECM and RNE-ECM, different laminin topologies could nevertheless result in local changes in mechanical properties. To explore this idea initially theoretically, we modeled the laminin matrix as a hexagonal spring lattice with a fixed boundary (*29*) (see Materials and Methods for a detailed description of the model). The topology of the matrix was then varied by changing the degrees of porosity in the lattice. Mapping the distribution of local tensions in the network revealed that with an increase in porosity, regions with higher tensions appeared (Fig. 4, A to B). In addition, using a simple theoretical model of a string under tension, we found that it should be harder to deform the string in the orthogonal direction as the tension in the string increases (Fig. 4, C to D). To test this idea experimentally, we made use of the fact that mitotic cells in the optic cup deform the underlying laminin when rounding apically (Fig 4,C). Analyzing laminin deformations indued by these mitotic cells, we found that in accordance the model they induce smaller deformations in the porous rim regions compared to the RNE region where laminin is less porous (Fig. 4, E to F and movie S7).

**Fig. 4.**
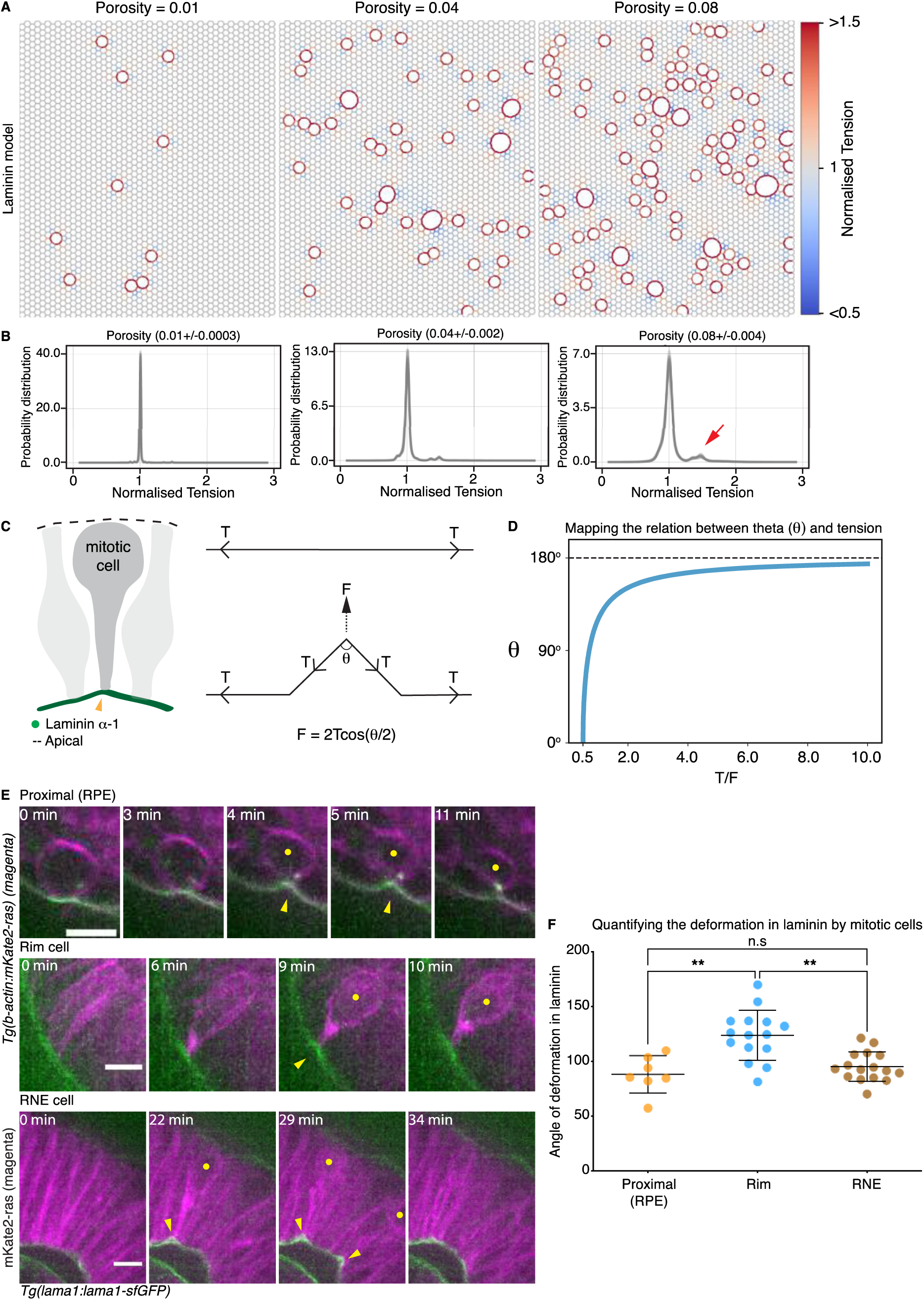
Changing topologies can influence the mechanical properties of laminin. (**A-B**) Laminin modeled as a hexagonal lattice with a three-point vertex with increasing porosity. Color bar indicates values of normalized tension (tension in porous lattices /tension in a non-porous lattice). Porosity is defined as the ratio of (number of nodes removed/total number of nodes in the network). The red arrow points to the appearance of springs under high tensions in the lattice on increasing porosity (n = 10 simulations) **(C-D)** Analysis of a perpendicular force on a string under tension. Graph depicts the relationship between the angle θ and the ratio of T/F (Tension in the string / Perpendicular force on string). Schematic of mitotic cell, yellow arrow points to deformation in laminin. **(E)** Montage of laminin deformations (green) on apical cell division in live *Tg(lama1:lama1-sfGFP)* embryos labeled with mKate2-ras (magenta) (scale bar = 10 μm). **(F)** Graph of the angle of deformation (mean with SD) in laminin on apical cell division in the proximal (n = 7 cells), rim (n = 15 cells) and RNE (n = 15 cells) regions in live *Tg(lama1:lama1-sfGFP)* embryos. Kruskal Wallis test with Dunn’s multiple comparison (** p = 0.025 for rim vs RNE, ** p = 0.029 for rim vs Proximal).

Thus, our theoretical model and experiments indicate that network topology can influence the distribution of local stresses within the network resulting in changing mechanical properties of the lamina.

### Focal adhesion dynamics and rim cell migration are less directed upon interference with laminin topology

Our theoretical model predicted that after a certain threshold of node removal and increasing porosity, large circular breaks should appear that distort the topology of the network nearby (fig. S4A). To test this prediction experimentally, the laminin structure was disrupted using a verified laminin α-1 morpholino (*18, 30*) to downregulate laminin. Morpholino treatment was preferred over a full mutant as it allowed the creation of varying phenotypes from mild disturbance to full ECM rupture, depending on morpholino concentration.

Embryos injected with 0.4ng/embryo of laminin α-1 MO into the *Tg(lama1:lama1-sfGFP)* or *Tg(lama1:lama1-mKate2)* lines showed severe laminin downregulation resulting in large laminin pores and breaks in the structure that were usually more severe in the rim region than the RNE (Fig. 5A yellow asterisks). Interestingly, the shape of the breaks and the topology of the surrounding laminin, strongly resembled the distorted topology observed on drastically increasing porosity in the theoretical lattice (fig. S5A).

**Fig. 5.**
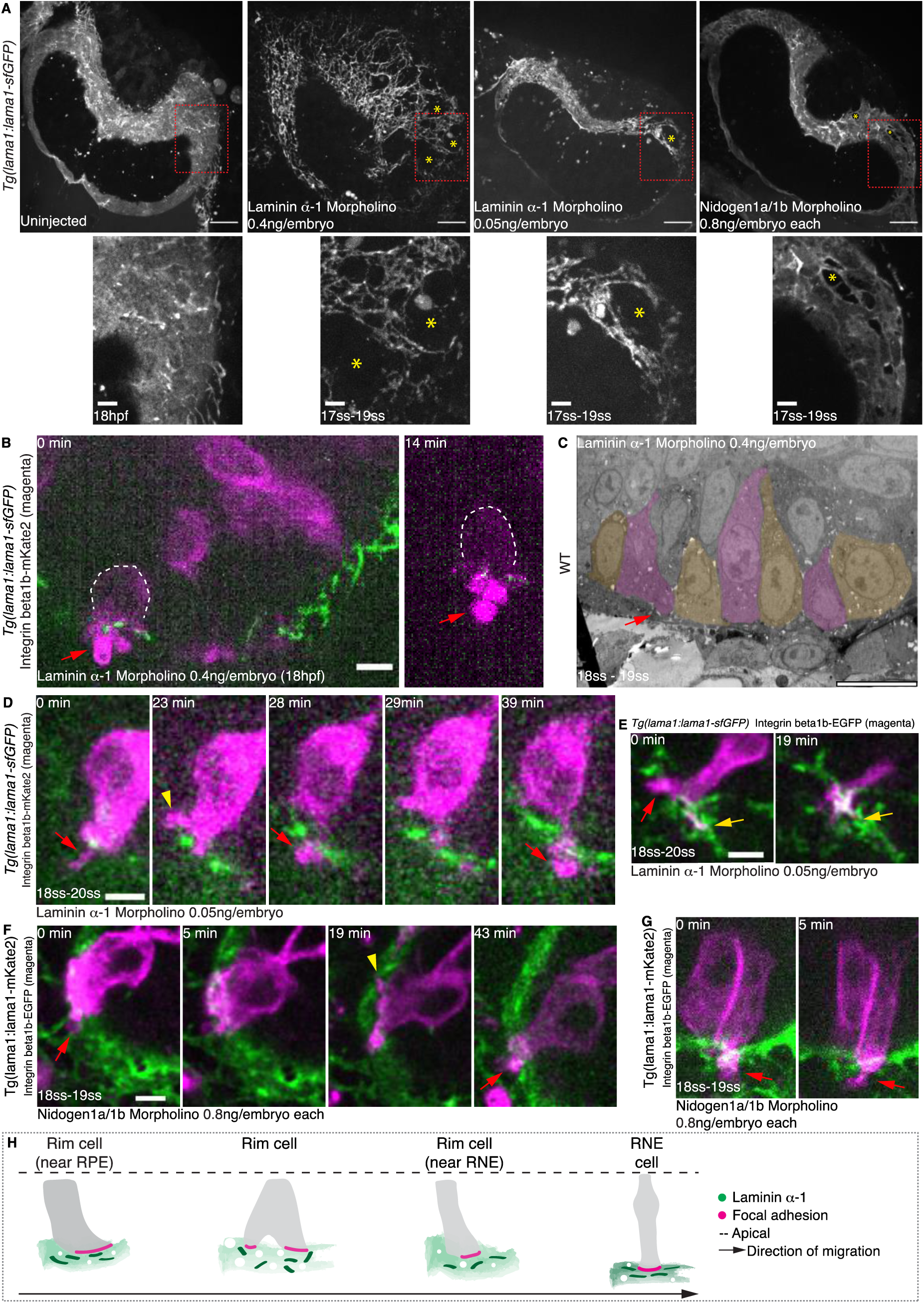
Interference with laminin topology leads to less directed focal adhesion dynamics and cryptopodia loss. (**A**) Maximum intensity projections of live *Tg(lama1:lama1-sfGFP)* embryos (scale bar = 20 μm) treated with 0.4ng/embryo of laminin α-1 morpholino (MO). Red dotted box indicates the region in the inlay (scale bar = 5 μm). Yellow asterisks indicate the breaks in the laminin structure. (**B**) Snapshot of live *Tg(lama1:lama1-sfGFP)* (green) embryo treated with 0.4ng/embryo laminin α-1 MO, with a rim cell labeled with Integrin beta1b- mKate2 (magenta). The red arrow points to a bleb. White dashed line outlines the cell (scale bar = 5 μm). (**C**) TEM image of WT embryo treated with 0.4ng/embryo of lama α-1 MO (scale bar = 10 μm). Red arrow points to a bleb like structure. Yellow and pink areas delineate single cells. (**D-E**) Montages of rim cells in *Tg(lama1:lama1-sfGFP)* (green) embryos treated with 0.05ng/embryo laminin α-1 MO and labeled with Integrin beta1b-mKate2 (magenta) in (scale bar = 5 μm). Red arrows points to a bleb. Yellow arrowhead points to a less directed FA developing in direction opposite to that of migration. Yellow arrows point to retracting leading FA. (**F-G**) Montage of rim cells in *Tg(lama1:lama1-mKate2)* (green) treated with 0.8ng/embryo Nidogen1a/1b MO labeled with Integrin beta1b-EGFP (magenta). Red arrows point to a bleb. Yellow arrowhead points to a less directed FA developing in direction opposite to that of migration (scale bar = 5 μm). **(H)** Schematic summarizing the different cell-matrix interactions observed on different topologies of laminin.

Next, we aimed to test whether disrupting the laminin topology influenced cell behavior. We imaged the FA dynamics along with laminin in laminin morphants and noted that rim cells started blebbing basally as reported previously (*18*) instead of throwing podia in the direction of movement. These blebs hindered migration and often extended through the laminin breaks (Fig. 5B). In most instances (7/9 cells) rim cells were stuck in the same position, unable to retract the blebs extending through the matrix (Fig. 5B, fig. S5D and movie S8). In regions where laminin structures were still more intact, cells did not get stuck but focal adhesions nevertheless exhibited less directed movements and cells migrated over shorter distances than in controls (fig. S5G). Lowering the concentration of the morpholino (0.05 ng/embryo) resulted in overall smaller breaks in the laminin structure (Fig. 5A) and close to equal porosity between the rim and RNE regions (fig. S5B). In this condition, rim cells blebbed but did not remain stuck. They nevertheless showed less directed FA movements and migration was overall less efficient (4/5 cells) (Fig. 5, D to E, fig. S5G and movie S8).

We additionally perturbed laminin arrangements without interfering directly with the protein’s expression by downregulating the matrix cross linker protein nidogen (*29*), known to be required for OCF (*31, 32*), using established morpholinos against nidogen 1a and nidogen 1b (see Materials and Methods). This condition phenocopied laminin knockdown and resulted in breaks with increased pore sizes in the laminin structure around the optic cup (Fig. 5A). We again observed rim cells blebbing through the laminin structure (Fig. 5, F to G and movie S9). Further, this condition led to cells with less directed focal adhesion dynamics (2/7cells) (Fig 5, F to G yellow arrow and fig. S5G). Milder perturbations in which the laminin matrix did not present as large breaks but was less organized, showed rim cells that migrated without blebs but stepped backwards more often than in controls (3/7 cells) (fig. S5G).

To understand how these conditions influenced the cells’ protrusive activity at the structural level, we imaged embryos injected with 0.4ng/embryo of laminin α-1 MO using TEM. We found that rim cells showed less or no cryptopodia-like protrusions (Fig. 5C). Instead, bleb-like structures were observed (Fig. 5C red arrow and fig. S5C), suggesting that disrupting the topology of laminin directly affects rim cell protrusion dynamics.

Our experiments show that downregulating laminin results in the disruption of laminin topology, confirming the prediction of the model. Additionally, we found that disrupting laminin topology leads to less efficient rim cell migration presenting a direct link between matrix topology and rim cell migration efficiency.

## Discussion

The use of new tools to label the ECM component laminin *in vivo* in combination with sophisticated image analysis pipelines enabled us to show that matrix topology guides collective cell migration *in vivo* during OCF. During zebrafish OCF, cell matrix interactions depend on basal cryptopodia and their dynamics change with differing matrix topology. Changing this topology upon genetic interference results in less directed migration. This and our photoconversion experiments reveal that rim cells are not dragged passively into the neuroepithelium as suggested previously (*19*) but actively migrate towards it. This active rim migration is driven by cryptopodia, which are defined as basal protrusions that extend beneath the next cell along the direction of migration (*22*). These have so far mainly been reported *in vitro* (*22, 23*) but recently they have also been suspected to be involved in some types of collective cell migration *in vivo* (*33, 34*). In ‘closed’ epithelial structures *in vitro* that lack a free edge or clearly defined leader and follower cells, the direction and angle of contact of cryptopodia with the underlying substrate predetermines the overall direction of migration (*23*). A similar mechanism could operate in the optic cup which resembles a closed epithelium, in which cryptopodia could act to break symmetry and define the direction of migration. The finding that FA movements over the matrix are seen as early as 13hpf before the onset of RNE invagination (Fig. 1L and fig. S1I), strengthens this idea. Interestingly, at these timepoints laminin is already differentially enriched in the RNE region (fig. S3G). However, the hierarchies between the timing of cryptopodia polarization, changes in matrix topology and rim migration onset need to be further explored, taking other cell behaviors that occur during OCF into account (*35*).

Rim cells migrate over changing laminin topologies, with laminin porosity being the highest in the center of the migration path which corresponds to the rim region (see schematic in Fig. 5H). This strongly suggests that rim migration is driven by topotaxis, a phenomenon not yet widely reported *in vivo*, where cells can migrate toward denser or sparser topological features (*17*). Since topotaxis is regulated by the intrinsic stiffness of the migrating cell, it will be important to understand how cell mechanics and matrix topology influence each other during OCF. We observed that cells on very large laminin breaks are unable to maintain their shape and display extremely ‘floppy’ behavior (fig. S5, E to F and movie S10), possibly indicating that they become ‘softer’. This implies a link between cell stiffness and matrix topology as shown *in vitro* (*17*). Investigating the relation and interplay between matrix topology and physical properties of the cell will be exciting areas to study in the future, comparing *in vitro* and *in vivo* data.

It will further be interesting to probe how cell-matrix interactions themselves could reorganize the matrix. *In vitro* it has been shown that endothelial cells rearrange the matrix to mimic the overlying branched EC network (*36*) and in the *Drosophila* egg chamber migrating follicle cells influence the progressive alignment of matrix fibrils (*8*). We also observe that during OCF, laminin becomes progressively aligned in the RNE where tissue invagination occurs (fig. S3, I to J), indicating a role for cell-matrix interactions in matrix arrangement.

Importantly, in addition to the developmental context, the arrangement of matrix molecules is crucial to wound healing and is disrupted in diseased states like cancer (*17*) and fibrosis (*37*). Thus, understanding the interplay between matrix topology, cell behavior and the factors that regulate ECM arrangement during development will facilitate the understanding of factors underlying diverse severe disease states. Such findings will also help create better *in vitro* environments for organoids, which are emerging as useful tools to understand human development and regeneration.

## Acknowledgements

We thank the Fish facility, LMF facility, Scientific Computing facility and the Transgenomics facility at MPI-CBG. Thanks to A.L. Sousa, and E.M. Tranfield from the Electron Microscopy Facility at IGC for TEM protocol design and sample processing and cutting. Noreen Walker is tanked for contributing the image rotation script used in the analysis and Gayathri Nadar for help and advice with image analysis. We thank Aleksandra Syta and Mihail Sarov for help with the development of BAC tagging lama1 to sfGFP, mKate2, dendra, Sylvia Kaufmann and Heike Hollak for technical help in screening the laminin α-1 transgenic lines and Elisa Nerli for technical help with the transplantation experiments. We thank Tiago Paixao from the IGC quantitative and digital science unit for advice and help calculating the statistics of significance for our quantifications. We thank the Norden lab and Modes lab members for critical comments on the project and manuscript. Thanks to Ivo Telley and Julien Vermot for critical remarks and review of the manuscript.

## Author contributions

Conceptualization: KGS, CN, CM

Methodology: KGS, APR, JS, CN, CM, JG, IFS, AS, KBH, AK, RS

Investigation: KGS, APR, JS, AK, AS, KBH, RS

Funding acquisition: CN, CM, JG, IZ

Supervision: CN, CM, JG, IFS

Writing – original draft: KGS, CN

Writing – review and editing: KGS, CN, CM, IFS, APR, JS, AK, AS, KBH, RS

## Materials and Methods

### Zebrafish husbandry

All fish were treated according to the European Union directive 2010/63/EU and the German Animal Welfare Act. Embryos were incubated at 28°C, until they reached the shield stage of development. Embryos were then transferred to 21°C until 12hpf (5ss-6ss) and further incubated at 32°C and 28°C in E3 medium to obtain stages between 7ss - 20ss (somite stage). All live-imaging was carried out at 28°C.

### Mounting and staging of zebrafish embryos for imaging

Zebrafish embryos were mounted with their heads facing the glass in MatTek glass bottom microwell P35G-1.5-14-C imaging dishes. Embryos were anaesthetized with 0.28 - 0.4μg/μl of MS-222 before imaging and mounted in 100μl of 0.6% ultrapure low melting point (LMP) agarose (Invitrogen REF 16500-500). Fixed and immunostained embryos were mounted in 100μl of 1% ultrapure LMP agarose. E3 medium was added to the imaging dish after mounting. The temperature was maintained using an objective heater or a temperature incubator.

Zebrafish embryos were staged according to Kimmel et al 1995 (*38*) by counting the somites before mounting. The following somite counts were used to stage the embryos at 13hpf (7ss-9ss), 14hpf (10ss-11ss), 16hpf (13ss-14ss), 18hpf (17ss-18ss), 19hpf (19ss-20ss) and 20hpf (21ss-22ss). Experiments with embryos covering different somite counts than those mentioned here are specified accordingly in the figure legends. The number of repetitions done for each experiment is documented in Table S6.

### Zebrafish lines and strains

TL Dresden+/-,TL Ultrecht+/-, *Tg(lama1:lama1-sfGFP)*, *Tg(lama1:lama1-mKate2)*, *Tg(lama1:lama1-dendra)*, *Tg(Vsx2:GFP)* (*39*), *Tg(ubi:ssNcan-EGFP)* (*40*) and Tg(β- actin:mKate2-ras) (*18*).

### Generation of the BACs tagging laminin to fluorescent proteins

BAC recombineering was performed in the transgenomics facility at MPI-CBG using a previously published protocol (*41*) (Protocol description adapted from Dr. Jaydeep Sidhaye thesis: https://nbn-resolving.org/urn:nbn:de:bsz:14-qucosa-232445). The BAC clones containing the *lama1* gene (CH211-170M3) were purchased from Source Bioscience. These BACs contained the pTARBAC2.1 backbone. Gene sequences were confirmed through PCR and sequencing (Table 1). BAC clones were first transformed with pSC101-BAD-gbaA-tet-plasmid which expresses the lRed operon proteins upon induction with the L-arabinose enzyme. The plasmid also contains a temperature sensitive replicon pSC101 and tetracycline (tet) selection marker.

Tol2 arms were integrated into the BAC clones using a Tol2-amp plasmid (obtained from Tatjama Sauka-Spengler (*42*)) featuring a R6K origin site incorporated into it that was cloned from a R6K-TARBAC-Tol2 plasmid (kind gift from Nadine Vastenhouw).

*E.coli* carrying the BAC clones and the pSC101-BAD-gbaA-tet-plasmid were grown in LB medium along with chloramphenicol and ampicillin at 3°C. The expression of λRed proteins was stimulated with L-Arabinose (0.2%) and the reaction was kept on a shaker at 37°C for 45 minutes. *E.coli* cells where then electroporated with the Tol2-amp plasmid and grown at 30°C. Positive colonies were selected for successful Tol2-amp integration and confirmed through sequencing (Table S1).

The *lama1* gene was tagged on the c-terminal with fluorescent proteins super folded GFP (sf-GFP), mKate2 and the photoconvertible dye dendra (Table S2). These tags contained a kanamycin selection marker that was flanked by FRT sites. These FRT sites can be removed by site specific Flp/FRT recombination. R6K plasmids carrying the respective tags were used to amplify 50bp of 5’ and 3’ homology arms to the *lama1* gene using specific oligos (Table S3). In order to tag the *lama1* gene with the different fluorescent proteins, *E.coli* cells carrying the BAC-Tol2-Amp clones along with the pSC101-BAD-gbaA-tet-plasmid were electroporated with the respective tagging cassettes and grown at 37°C. At this temperature the pSC101-BAD-gbaA-tet-plasmid is not replicated and gets slowly removed from the culture. The kanamycin marker from the fluorescent tags were removed by electroporating the cells with pRedFlp-hgr plasmid. This plasmid contains anhydrotetracyclin dependent Flippase (Flp) expression, a temperature sensitive replicon pSC101 and the selection marker hygromycin.

*E.coli* cells now containing the tagged BAC-Tol2-Amp clones and the pRedFLP-hgr plasmids were grown again in an LB medium that contained the antibiotics chloramphenicol, ampicillin, hygromycin along with the enzyme anhydrotetracyclin (200ng/ml) for 3 hours at 30°C (see Table S4 for antibiotic concentrations). Cells were grown on plates with and without kanamycin to test for sensitivity towards the antibiotic. *E.coli* colonies that only grew on plates that lacked kanamycin were selected.

### Zebrafish transgenesis

The BAC plasmid was injected at 0.1ng/embryo along with 0.3 - 0.6ng/embryo of transposase2 enzyme mRNA into the cell of 1 cell staged WT embryo. F0 embryos were screened at 24 hpf for fluorescence signal using a fluorescence stereoscope and positive embryos were selected and grown. F0 adults were outcrossed to WT fish and the progeny was screened for positive fluorescence signal and by PCR on the whole genome (Table S5). The enzyme emerald was used in the PCR reaction to amplify the GFP sequence in the whole genome sample. F0 progeny that expressed GFP were selected and grown. F1 adults were screened by tail fin clipping and checking the genome for GFP expression. Positive F1 fish were then outcrossed to WT fish and the progeny was further screened on the confocal microscope for GFP signal at 24hpf. The F1 with the highest frequency of positive progeny and with the strongest fluorescence signal was used to further propagate the line. Subsequent generations were screened for fluorescence signal at 24hpf using a fluorescence stereoscope. This procedure was used to screen and propagate the *Tg(lama1:lama1-sfGFP)*, *Tg(lama1:lama1-mKate2)* and *Tg(lama1:lama-dendra)* lines.

### Transplantations

Donor embryos *Tg(lama1:lama1-sfGFP)* were injected with H2B-RFP mRNA to label the nucleus of donor cells. Donor and acceptor embryos were dechorionated with pronase at the 1000 cell stage and washed with Daniaeus buffer solution before transplantation. About 50 donor cells were transplanted into the animal pole of the acceptor embryos *Tg(lama1:lama1-mKate2)* at high stage of zebrafish development. Transplanted embryos were then incubated at 32°C until shield stage and transferred to a glass dish for further incubation at 21°C with 100 units of penicillin and streptomycin. Embryos were mounted in agarose and imaged at 16ss - 19ss.

### mRNA injections

Zebrafish embryos were injected with 5-40pg mRNA transcribed from the plasmids pCS2+ Integrin beta1b-EGFP, pCS2+ Integrin beta1b-mKate2 (*18*), pCS2+ Paxillin-mKate2 (*18*) and PCS2+ mKate2-ras (*43*) at 64 - 128 cell stages to obtain mosaically labeled FAs in cells in the optic cup. Transcription of mRNA was done using the Ambion mMessage mMachine kit.

### Morpholino injections

Morpholinos were injected into the yolk of the embryo at the one cell stage at concentrations of 0.05 – 0.4 ng/embryo of laminin a-1 MO 5’-TCATCCTCATCTCCATCATCGCTCA-3’ (*30*). To downregulate the ECM crosslinking protein nidogen, morpholinos against nidogen1a 5’- GTGCCGACCCATATCCAGTCCCAAA-3’ and nidogen1b 5’- CGGCATCTTCCCCAGGTAGTCAGAC-3’ were co-injected at a concentration of 0.8 ng/embryo each (*31*) into the yolk of the zebrafish embryo at the single cell stage.

### Immunostaining

Embryos were dechorionated at the required stages and fixed overnight at 4°C in 4% PFA solution on a rocker. Fixed embryos were washed 5 times with PBT (PBS + 0.8% Triton X-100) for 10 minutes and then put on ice and treated with trypsin for 15 minutes. This was followed by on ice incubation with PBT for 30 min. Embryos were then washed with PBT for 5 minutes on a rocker at room temperature. Fixed and permeabilized embryos were incubated overnight in 100 μl of 1:10 (diluted with PBT) NGS (normal goat serum) blocking solution at 4°C on a rocker. Fixed, permeabilized and blocked embryos were incubated with 100ml of primary antibody at the respective concentrations (diluted with PBT) and 1:100 NGS blocking solution for 48 hours at 4^0^C on a rocker. This incubation was followed by 5 PBT washes for 30 minutes each. Embryos were then incubated with 100 μl of secondary antibody at the respective concentration (diluted with PBT) and 1:100 NGS blocking solution for 48 hours at 4°C on a rocker. Embryos were then washed 4 times with PBT for 15 minutes each and afterwards 2 times with PBS for 10 minutes each on a rocker. Stained embryos were stored at 4°C in PBS.

Primary antibodies were used at the following concentrations: 1:100 anti-laminin (laminin α-1, RRID:AB_477163, Sigma L-9393), 1:100 anti-GFP (RRID:AB_94936, Milipore MAB3580) Secondary antibodies were used at the following concentrations: 1:500 Alexa Fluor 568 donkey anti-rabbit (RRID:AB_2534017, Thermo Fischer Scientific A10042), 1:500 Alexa Flour 488 chicken anti-mouse (RRID:AB_141606, Invitrogen A21200), DAPI (1:10,000).

### Characterizing the new laminin transgenic lines

*Tg(lama1:lama1-sfGFP)* zebrafish embryos fixed and stained at 24hpf, with anti-GFP and anti-laminin antibodies (laminin α-1), were imaged on the LSM 880 inverted confocal microscope with a Plan-Apochromat 20X (0.45NA) M27 air objective.

### Fluorescence recovery after photobleaching

FRAP experiments were done on an Andor spinning disk FRAPPA with the iXON 897 monochrome EMCCD camera using an Olympus UPLSAPO 30x (1.05NA) silicon oil objective. Region of interests (ROIs) were specified, bleached and imaged every 50ms for 5 minutes in a single xy plane. The ROI were imaged before bleaching for 20 time points at an interval of 50ms. To quantify the dynamics of laminin turnover, larger ROIs of bleaching were specified within the RNE and rim region and the imaging was done over a volume of 34.3μm (step size 0.7μm) in the axial direction, with a temporal resolution of 3 minutes for 1 -2 hours. The ROI was imaged for 20 time points at temporal resolution of 300ms before bleaching.

### Imaging cell-matrix interactions

Image acquisition was carried out on an Andor IX 83 inverted stand spinning disk and an Andor IX 71, inverted stand spinning disk. Olympus UPLSAPO 40x (1.25NA) and Olympus UPLSAPO 60x (1.3NA) silicon oil immersion objectives were used. The Andor IX 71 carried a Neo sCMOS camera and the Andor IX 83 spinning disk an Andor iXon Ultra 888, monochrome EMCCD camera. *Tg(lama1:lama1-sfGFP)* embryos were injected with Integrin beta1b-mKate2 mRNA and *Tg(lama1:lama1-mKate2)* embryos were injected with Integrin beta1b-EGFP mRNA and mounted at 17ss - 19ss to image interaction of FAs with laminin and tracked over at least 30 minutes with a temporal resolution of 0.58 - 1 minute.

For movies of morphants the temporal resolution varied from 1 - 5 minutes to avoid bleaching of the already weak laminin fluorescence signal and FAs were tracked for at least 30 minutes.

### Tracking the movement of FA relative to movement of the ECM

All photoconversion experiments were done on an Andor IX 83 inverted stand spinning disk with an Olympus UPLSAPO 60x (1.3NA) silicon oil objective. *Tg(lama1:lama1-dendra)* embryos were injected with Integrin beta1b-mKate2 at 64 - 128 cell stage to mosaically label focal adhesions in cells within the developing optic cup. Isolated rim cells with labeled FA were selected and a small region of the matrix beneath the cell was photoconverted over 5 time points at an interval of 1 minute. The movement of the FA was then tracked for at least 30 minutes at 1-minute intervals.

### Measuring the physical properties of the matrix using Brillouin microscopy

Live *Tg(ubi:ssNcan-EGFP)* embryos at 17ss were transferred to a glass bottom dish and incubated with 1 drop of 0.01% and 0.02% tricaine for 1 minute. The embryo was then mounted in 1% low gelling agarose and placed on ice for 1 minute. Imaging was done using a Zeiss C-Apochromat water immersion 40X (1.2 NA) objective. Snapshots were taken with transmitted light to document the region in which the measurement was done. Measurements were done on 5μm X 15μm ROI in the invagination and rim region. For the rim region, the ROI was placed at positions at which the optic cup was closely opposed to the overlying epidermis to avoid the signal from the surrounding yolk.

A previously published Brillouin set was used (*27*). In brief, the setup consists of a frequency-stabilized diode laser with a wavelength of 780.24 nm (DLC TA PRO 780; Toptica, Gräfelfing, Germany) and a two-stage virtually imaged phased array spectrometer. Adding a Fabry-Pérot interferometer (FPI) suppressed strong reflections from the glass surface. The data acquired was evaluated using a custom MATLAB program (https://github.com/BrillouinMicroscopy/BrillouinEvaluation).

### Transmission electron micrography workflow

Wild type (WT) embryos and WT embryos injected with lama1 morpholino were fixed in 2% formaldehyde (Electron Microscopy Science #15710), 2.5% glutaraldehyde (EMS #16220) and 0.1M Phosphate Buffer (PB) overnight at 4°C. To ensure complete penetration of the fixative, samples were immersed in fresh fixative and further processed in a PELCO Biowave Pro+ Microwave (Ted Pella), in a series of 7 cycles of 2 minutes each with an alternating irradiation power of 100W or 0W, followed by washes in0.1M PB. Samples were then post-fixed in a solution 1% osmium tetroxide (EMS #19110), 1% potassium ferrocyanide (Sigma-Aldrich #P3289-100G) in 0.1M PB in the same irradiation power cycles, followed by washes in 0.1M PB and milliQ Water. Staining was performed with 1% aqueous tannic acid (EMS #21700) followed by 1% aqueous uranyl acetate (Analar #10288) (7 cycles of 1 minute each with an alternating irradiation power of 150W or 0W). Samples were dehydrated in a graduated series of ethanol (VWR #VWRC20821.330) using a cycle of 40 seconds each with 150W power. Embedding was done using EMbed-812 epoxy resin (EMS #14120) after an infiltration series of 25%, 50%, 75% and 100% (each using a 3 minute cycle with 250W power), followed by polymerization at 60°C overnight

Sections of 70nm were cut using an ultra45 diamond knife (Diatome) on a UC7 Ultramicrotrome (Leica) and collected on slot grids coated with 1% formvar in chloroform. The sections were post-stained sequentially with uranyl acetate and lead citratefor 5 minutes each and then imaged on a Tecnai G^2^ Spirit BioTWIN Transmission Electron Microscope (TEM) from FEI operating at 120 keV. Samples where recorded with an Olympus-SIS Veleta CCD Camera using SerialEM software (*44*).

### Documenting laminin topology

Qualitative analysis of laminin topology was done with fixed zebrafish embryos of the transgenic lines *Tg(lama1:lama1-sfGFP)* stained with anti-laminin and anti-GFP antibodies. Embryos were imaged using a Zeiss LSM 700 inverted confocal microscope with pixel size 0.223 μm:0.223 μm:0.7 μm (x:y:z).

Fixed zebrafish embryos of transgenic line *Tg(Vsx2:GFP)* over stages of 18hpf and 19hpf were stained with anti-laminin (laminin α-1) and anti-GFP antibody. Embryos were imaged with a LSM 880 inverted confocal microscope with pixel size 0.92 μm:0.92 μm:0.1 μm (x:y:z). The Zeiss LD LCI planApochromat multi-immersion 40X (1.2NA) objective was used. Oversampling in the axial direction was done to facilitate the deconvolution of the images using the Huygens deconvolution software.

In both the experiments, laser power correction was used along with spline interpolation in the axial direction.

Morpholino treated *Tg(lama1:lama1-sfGFP)* embryos were mounted without fixation at 17ss - 19ss and snapshots were taken on the Andor IX 83 inverted stand spinning disk with Olympus UPLSAPO 60x (1.3NA) silicon oil objective. This was done to qualify and quantify the breaks in the laminin structure upon treatment with laminin α-1, nidogen1a and nidogen1b morpholino.

### Documenting the deformation of laminin on apical cell division

Deformations in laminin on apical cell division was observed in *Tg(lama1:lama1-sfGFP)* embryos injected at the single cell stage with mKate2-ras mRNA or crossed to the *Tg(β-actin:mKate2-ras)* transgenic line in order to visualize cells in the optic cup. Embryos were mounted at 14ss – 18ss and imaged for at least an hour at an interval of 0.58 minutes to capture the deformation of laminin on apical cell division. Image acquisition was carried out on an Andor IX 71, inverted stand spinning disk with the Neo sCMOS camera. Olympus UPLSAPO 40x (1.25NA) silicon oil immersion objective was used.

### Fluorescence recovery after photobleaching analysis

The same ROIs as specified during photobleaching were used to track the recovery of fluorescence. For the analysis, time points during the bleaching event were not considered for all the embryos in the rim and RNE regions. The fluorescence recovery intensity was normalized against the noise in the image and against acquisition bleaching (*45*). The ROI to correct for acquisition bleaching was specified in the ventricle where a stable laminin signal was observed. The position of the ROI was corrected in case of drift in the image. Normalized fluorescence recovery intensity was plotted for 3 minutes after bleaching (Fig. 1, G to H and fig. S1D).

### Quantifying laminin turnover (hours)

An ROI of 19×21 pixels was specified in the center of the photobleached region in the rim and RNE region (see fig. S1E). Imaging was done over 34.3 μm (step size 0.7 μm) and the intensity of fluorescence recovery was measured over a volume of 3.5 - 4.9 μm within the rim and RNE region covering 2-3 slices above and below the bleached region over 1 - 2 hours. The intensity was normalized to eliminate background noise and acquisition bleaching as mentioned above.

### Tracking and quantifying focal adhesion (FA) dynamics

A novel image analysis pipeline was used to segment the FA sites on curved surfaces (here: laminin structure) (*24*). We created a binary mask using the watershed algorithm to select the membrane with FAs to be segmented and to eliminate additional membranes before applying the pipeline. Each movie was treated separately with parameters fine-tuned for optimal segmentation of the FA. The centroid of the segmented FA was then tracked (*46*) over the length of the movie for at least 30 minutes generating a 3D track over time. In the event of the FA splitting or a new FA forming in the same cell, as seen for cells in the rim region, the FA that interacted with the laminin the longest was tracked to avoid interpolating the FA position. Drift in the movies was corrected by subtracting the velocity of the centroid of the ECM surface (marked as a stable point) from the centroid of the segmented FA.

3D tracks of segmented FAs were processed using the published MATLAB pipeline MSDanalyzer (*47*) (https://github.com/tinevez/msdanalyzer) to calculate the time averaged mean square displacement (tamsd) and the velocity autocorrelation. To extract the tamsd of a single FA track, overlapping sub-tracks of 20 minutes were created from the track and then analyzed with the MSDanalyzer. The tamsd was then calculated for each sub track for every cell. The velocity autocorrelation was computed at 30 time points of the FA tracks without creating sub tracks.

### Segmentation of laminin pores

Deconvolution was done in the Huygens deconvolution software. The backprojected pinhole radius (259.82 nm) was calculated using the automatic calculator available on the Huygens software inputting the pinhole side (35.1 μm) and the specification of the microscope LSM 880. To select the noise value, a line is drawn across the laminin signal and the lowest value of noise was selected. Other parameters set are the signal to noise ratio of 30, 50 iterations and a quality threshold of 0.05. The deconvolved file was the saved as a 16-bit file.

The pores within the laminin were segmented using the Weka segmentation tool in Fiji. A classifier was created by training on the data and this classifier was then used to process all embryos. ROIs of 200×200 pixels were projected 1.0 μm thick in the different regions of the optic cup. The area of the pores was calculated within the ROI outlining the laminin signal. The range for the particle size to be analyzed was set to 0-20 μm^2^ as few pores larger than 20 μm^2^ were found. These parameters were kept for all the different regions of the optic cup analyzed. Percentage porosity was calculated using the following formulae:

[Percentage porosity in projected area of laminin = Total area of pores / (Total area of pores + Total are of laminin signal) * 100]

To measure the projected porosity (%) in embryos treated with 0.05ng laminin α-1 MO, a separate classifier was created. ROIs of 200×200 pixels were created in the rim and RNE region and projected 10.0 μm thick to capture the large breaks. The range of the particle sizes was set to 0-infinity μm^2^ in order to quantify all large breaks in both regions. The porosity was then calculated in the same manner as done for the control embryos.

### Quantifying the arrangement and coherency of laminin

The arrangement of laminin was quantified using the same fixed samples used to map the porosity. Deconvolved images were used, and ROIs of 200×200 pixels were projected 2.0 μm thick and rotated to the x-y plane using a novel image analysis pipeline. This was done to avoid artefacts created by a simple maximum projection that could influence the orientation analysis. A custom Fiji code written by Noreen Walker, Image Analysis facility, MPI-CBG (https://dx.doi.org/21.11101/0000-0007-F43E-1) fits a surface through the 3-dimensional ROI by a combination of rotation with fitting a minimal-cost z surface (*48*), and max-projects along its average normal vector to obtain a 2D ROI of isotropic resolution.

In the 2D ROIs, oriented structures were detected by the gradient of image intensity, computed with a Scharr operator of kernel size 5 pixels to minimize orientational biases in the estimation from pixelated grayscale images (*49*).

Nematic vectors 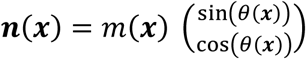 perpendicular to the gradient define object orientation. Thereby, *θ* and *θ* + 180° represent the same orientation or nematic vector. The prefactor *m(x)* allows to weight vectors for the subsequent analysis, and is chosen as the gradient’s magnitude. To avoid biases in orientations due to projections of the anisotropic point spread function, the original 3D stack was acquired with oversampling in z-direction [0.09 μm : 0.09 μm : 0.1 μm (x:y:z) resolution], and the overall distribution of orientations across each single ROI was checked to be isotropic.

The degree of local alignment between nematic vectors 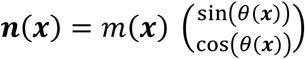 also known as coherency is defined within boxes of size 21×21 pixels from the ***Q*** -tensor

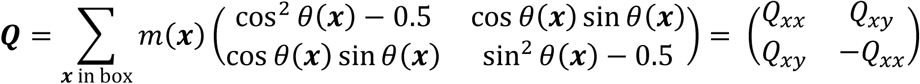

as twice its larger eigenvalue 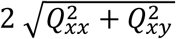, a quantity which is also known as scalar nematic order parameter. Averaging across all 21×21 boxes fully contained within the foreground, defined as the convex hull of the binarized image after smoothing with a Gaussian of kernel radius 5px, yields the mean local coherency (Fig. 3D and fig. S3J).

### Laminin intensity measurements

The mean intensity of laminin was measured in different regions of the optic cup over several microns in depth in live *Tg(lama1:lama1-sfGFP)* and *Tg(lama1:lama1-mKate2)* embryos at 17.9hpf (115 minutes from 16hpf). The mean intensity was sampled 14 μm from the dorsal most part of the optic cup to 49 – 56 μm in depth with an interval of 7 μm. The mean intensity of laminin in the different regions were normalized to the RNE region at each depth and was always found to be <1. These normalized intensities were then plotted and statistically analyzed (Fig. 3B).

### Laminin deformation measurements

The deformation observed in laminin on apical division in different regions of the optic cup was measured using the angle tool in Fiji. The quantification was done on maximum intensity projected time lapse images covering the entire region of laminin that was deformed by the dividing cell. These movies were sometimes corrected for noise by subtracting the background signal using a rolling ball radius of 500 pixels and adding a gaussian filter of sigma 0.2 μm radius to visualize the laminin better.

### Montage of time lapse images

Montage of time lapse images tracking FA dynamics in WT and morpholino conditions was done from maximum intensity projections of cropped movies that were manually registered and drift corrected using a Fiji plugin (http://imagej.net/Manual_drift_correction_plugin) developed by Benoit Lombardot (Scientific Computing Facility, MPI-CBG). In addition, the images were corrected for noise by subtracting the background signal using a rolling ball radius of 500 pixels and adding a gaussian filter of sigma 0.2 μm radius.

### Statistical analysis

Statistical analysis was done on GraphPad Prism 9 software. Confidence intervals of 95% and above were used. Detailed description of the tests and p values can be found in the respective figure legends. The table of replicates can be found in Table 6. To test the significance of the Brillouin shift between the rim and RNE across replicates, we used a linear mixed effects model, with a random intercept and region slope across replicates. The model was fitted using the python package statsmodels version 0.12.2.

### General description of the spring network model

#### Generating hexagonal lattice with pores

We model the laminin network as a hexagonal lattice of springs. For this we first generate a perfect hexagonal lattice consisting of 5200 vertices. Next, in order to generate pores, we randomly delete *n* vertices and their edges. We ensure that none of the vertices that are on the boundary are deleted. This leads to some vertices that have less than three neighbors (fig. S6A). We wish to generate a network where every vertex has only three neighbors, similar to the laminin network (*29*). Hence, we go through each remaining vertex and delete any vertex that has one or fewer neighbors. Vertices which have only two edges are deleted and the neighbors are connected if they are closer than twice the lattice edge length. Again, we do not delete any vertex that is on the boundary of the network. This algorithm gives us a lattice where every vertex that is not on the boundary has three neighbors (fig. S6B). We define the porosity of our lattice as the ratio between the number of vertices deleted and the number of vertices before generating pores (5200). For a given *n*, we generate ten networks using different random seeds. This gives us networks of very similar porosity values.

#### Implementing stress on the network

Each spring in the network has a spring constant, *k*, and a preferred rest length, *l_o_*. We keep the value of *l_o_* to be the same as the edge length of a perfect hexagonal lattice without pores. Also, the spring constant for all springs is given a value of 1. Next, we place the vertices of the network such that the whole network is uniformly stretched in all directions by a factor of s and fix the boundary vertices at their new positions. The rest of the vertices are then moved in discrete timesteps until a force balance is achieved.

In order to move the vertices, we assume overdamped dynamics which implies that the vertices have no inertia. After a given discrete timestep Δ*t*, the new position vector of a vertex *α*, *<u>x</u>^α^*, is given by

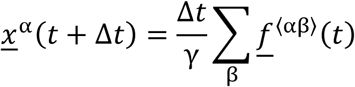

where *<u>f</u>*^(αβ)^ represents the force exerted by the spring connecting vertex α with its neighbor *β*. *γ* represents the friction coefficient and is kept as 1 arbitrarily. The force on the spring is computed as follows

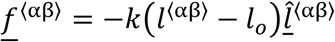

Where 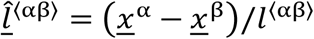 and 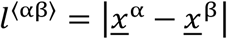.

#### Calculating the normalized tension in the springs

In order to quantify the amount of stretch and compression in the networks after achieving equilibrium, we compute for each spring a *normalized tension*. This is the ratio between the signed scalar force in springs and the expected force in springs of a perfect hexagonal lattice with no pores. The normalized tension for a spring connecting vertices *α* and *β* is given by

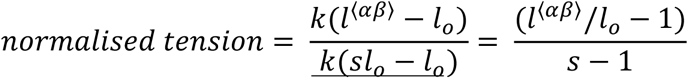

This quantity is a scalar dimensionless quantity computed for each spring. A value > 1 indicates springs stretched more than the stretch in springs in a full network without any pores. Similarly, value < 1 indicates relatively less stress than a typical spring in a full hexagonal network. The codes to perform the simulation are available here: https://git.mpi-cbg.de/krishna/laminin_network.

### Observations

We observe that on achieving force balance, the pores in the network take up circular shapes (fig. S7). Moreover, for networks with very high porosity, we see small pores elongating along the larger pores that look similar to the pores observed in the laminin structures in embryos treated with 0.05ng/embryo of laminin α-1 MO (fig. S5A).

We also observe that with increase in porosity up to an extent, we see more stretched springs appearing. This can be observed as a new peak in the distribution plots of the normalized tensions (fig. S8). We also observe a very slight but consistent increase in the mean normalized tensions with increase in porosity. However, on increasing the porosity value beyond 0.2, we start seeing a slight decrease in the mean normalized tension and the distribution starts to flatten out (fig. S9A). More stretched springs take up more space in the network than less stretched springs. In order to quantify the space of the network that ‘appears’ stretched to the cell, we quantified the length weighted normalized tension. Here again, we see a slight but consistent increase in the mean value with increase in porosity, except for networks with very high porosity (fig. S9B). It should be noted that these simulations with high porosity (>0.2) qualitatively look much more porous than the images of the laminin network in the optic cup in wild type conditions.

---

**Fig. S1.**
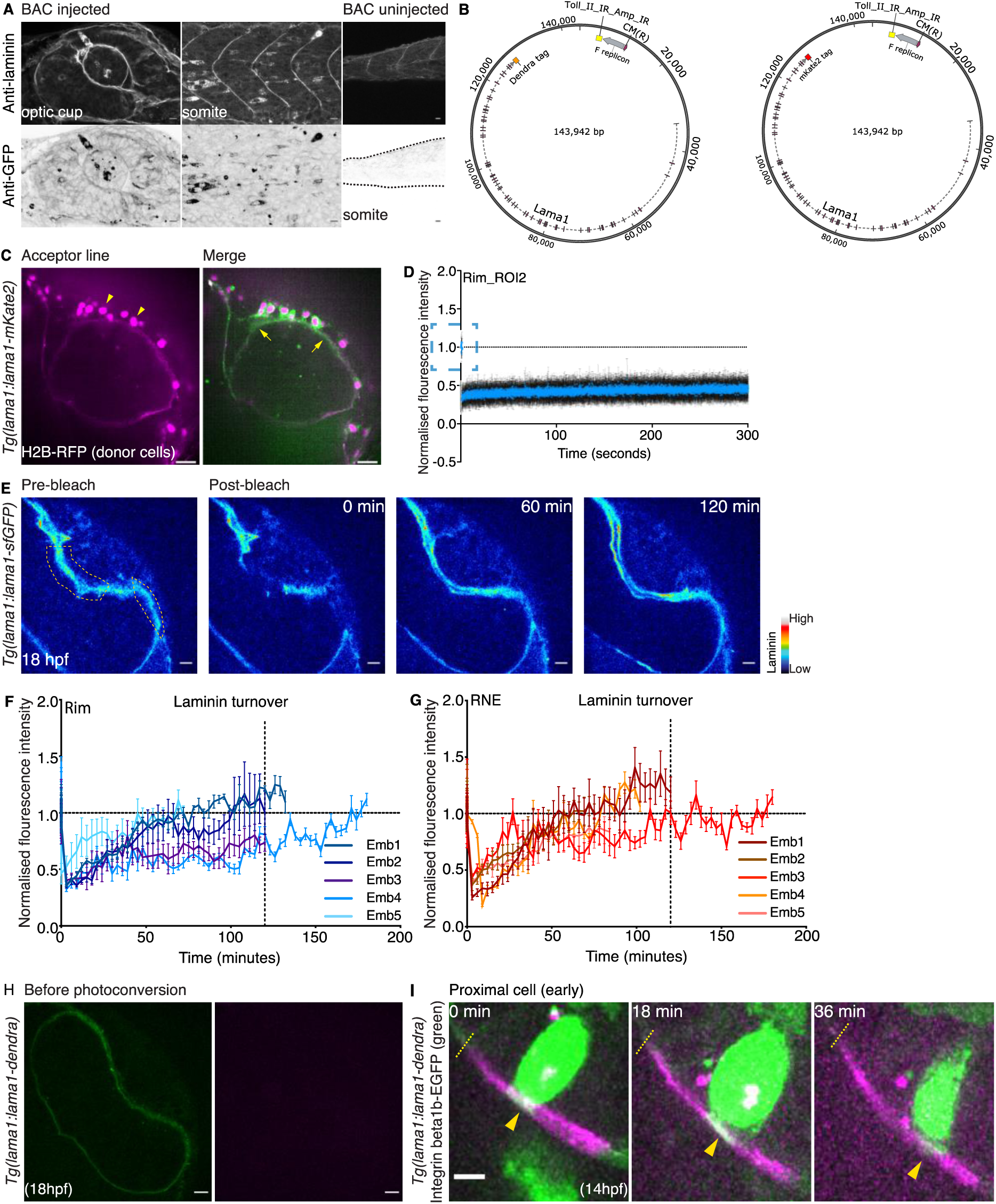
Laminin turns over within hours and can be deposited by tissues adjacent to the optic cup. **(A)** Maximum intensity projected snapshots of the optic cup and somite at 24 hpf of fixed WT embryos injected with lama1::lama1-sfGFP BAC (scale bar = 10 μm). Uninjected embryos at 24hpf are shown as maximum intensity projected snapshots of the somite of a fixed WT embryo. Embryos are stained against laminin α-1 (grey) and GFP (black) antibodies (Images modified from Dr. Jaydeep Sidhaye thesis: https://nbn-resolving.org/urn:nbn:de:bsz:14-qucosa-232445). **(B)** Schematic of zebrafish *lama1* BACs (CH211-170M3) tagged to the red fluorescent protein mKate2 and the photoconvertible dye dendra. **(C)** Maximum intensity snapshots of live *Tg(lama1:lama1-mKate2)* (magenta) acceptor embryos transplanted with cells from *Tg(lama1:lama1-sfGFP)* (green) donor embryos. Donor cells are tagged with nucleus marker H2B-RFP (magenta) to identify them (scale bar = 20 μm). **(D)** Normalized fluorescence intensity plot of laminin in the rim ROI1 region (mean and SD, n = 7 embryos). **(E)** Montage of live *Tg(lama1:lama1-sfGFP)* embryos (royal LUT) mounted at 18hpf pre and post bleaching ROIs in the rim and RNE region (scale bar = 10 μm). Dotted yellow shape outlines the bleached region. **(F-G)** Normalized fluorescence intensity plots of laminin in the rim and RNE region (mean and SD, n = 5 embryos per experiment). **(H)** Snapshot of live *Tg(lama1:lama1-dendra)* (green) embryo before photoconversion (scale bar = 10 μm). Magenta snapshot shows no signal before photoconversion. **(I)** Montage of live *Tg(lama1:lama1-dendra)* embryos in which laminin was photoconverted (magenta) and focal adhesions were labeled by Integrin beta1b-EGFP (green) (scale bar = 5 μm). Dashed yellow lines indicate the start of the photoconverted laminin region. Yellow arrowheads point to moving FAs.

**Fig. S2.**
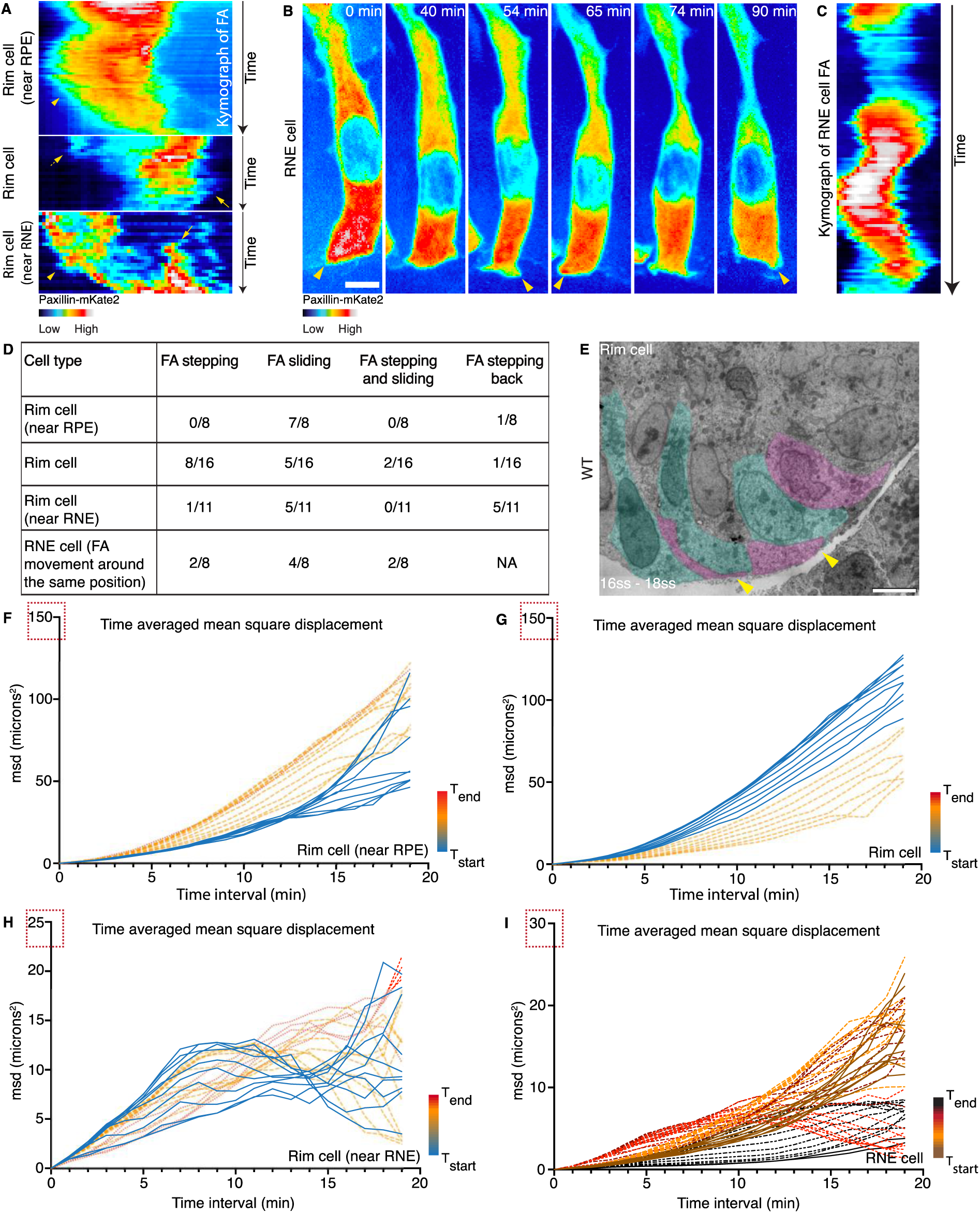
Rim cells near the RNE and RNE cells exhibit less directed FA movements. **(A)** Kymograph of FA of migrating rim cells near the RPE, in the center of the rim region and near the RNE. Yellow arrows point to newly forming leading FA, yellow arrowheads point to sliding FA. Dotted yellow arrow points to detaching FA. **(B)** Montage of non-migrating RNE cells. Focal adhesions are labeled with Paxilllin-mKate2 (royal LUT). Yellow arrowheads point to newly forming FA (scale bar = 5 μm). **(C)** Kymograph of FA of RNE cell (n = 3 cells). **(D)** Table documenting the frequency of different FA behaviors observed in migrating rim cells near the RPE, RNE, in the center of the rim and in non-migrating RNE cells. **(E)** TEM image of basal protrusion of cells in the rim region of WT embryos (scale bar = 5μm). Yellow arrowheads point to cryptopodia like protrusions in rim cells. Green and pink areas delineate single cells. **(F-I)** Time averaged mean square displacement (tamsd) plots of sub tracks of migrating rim cells near RPE and RNE, in the center of the rim and in the RNE region (means plotted). Color code indicates increasing time from the start to the end of the movie. Red boxes show the differences in maximum range of tamsd for migrating cells in different regions of the rim and non-migrating RNE cells.

**Fig. S3.**
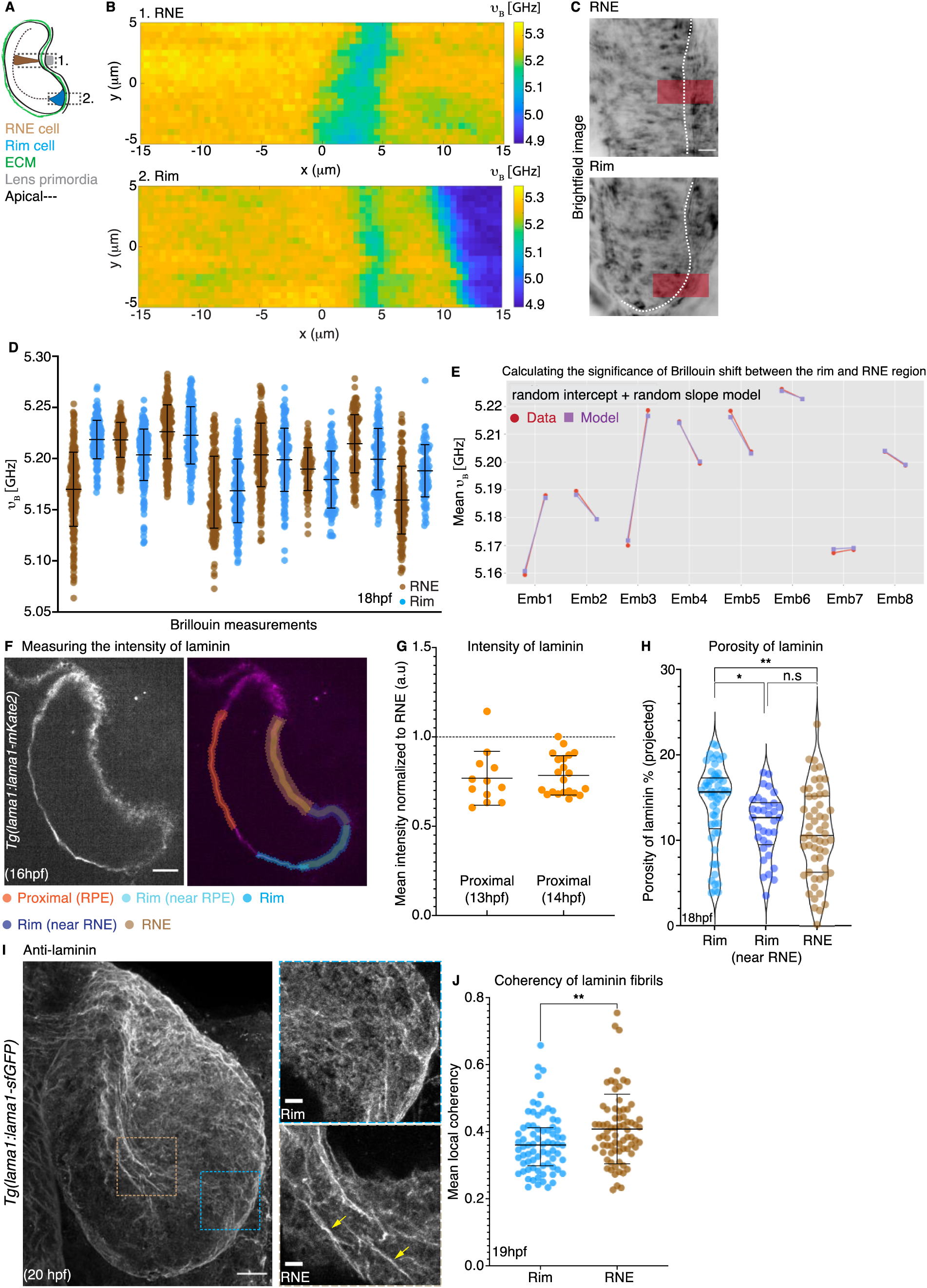
Matrix compressibility does not show a stark gradient while laminin topology differs at different stages of OCF. **(A)** Schematic of optic cup at 18hpf. Dashed black boxes outline the ROIs of Brillouin measurement. **(B)** Example of Brillouin measurement done in the rim and RNE region. **(C)** Transmitted light snapshots of live *Tg(ubi:ssNcan-EGFP)* embryos in the rim and RNE regions with red boxes roughly outlining the ROI of Brillouin measurement. White dotted line delineates the optic cup from the adjacent lens tissue (scale bar = 10 μm). **(D)** Graphs of Brillouin measurements in the rim and RNE regions (n = 8 embryos). **(E)** Graph mapping the random intercept and slope model to the experimental mean values of the Brillouin shift for each embryo. Using this model, the overall difference in means between the rim and RNE region was found to be insignificant (p = 0.636). **(F)** Snapshot of live *Tg(lama1:lama1-sfGFP)* embryo with color coded ROIs outlining the region of laminin intensity measurement (scale bar = 20 μm). **(G)** Intensity measurements of laminin in the proximal regions normalized to the mean intensity of laminin in the RNE region (means with SD) at 13hpf (n = 2 embryos each) and 14 hpf (n = 3 embryos each). **(H)** Graph of porosity of laminin (% projected) in the rim (n = 8 embryos), rim near RNE (n = 4 embryos) and the RNE (n = 8 embryos) regions. Kruskal-Wallis test with Dunn’s multiple comparison test (median with quartiles, * p = 0.0389, ** p = 0.0010). **(I)** Maximum intensity projection of fixed *Tg(lama1:lama1-sfGFP)* embryos at 20hpf stained with against laminin α-1 (grey). Brown and blue dotted boxes mark the region of RNE and rim in the inlay (scale bar = 20 µm). Yellow arrows point to laminin fibrils (scale bar = 5 µm). **(J)** Graph of mean local coherency of laminin fibrils at 20hpf (n = 4 embryos). Two tailed Mann Whitney test (median with interquartile range, ** p = 0.0069).

**Fig. S4.**
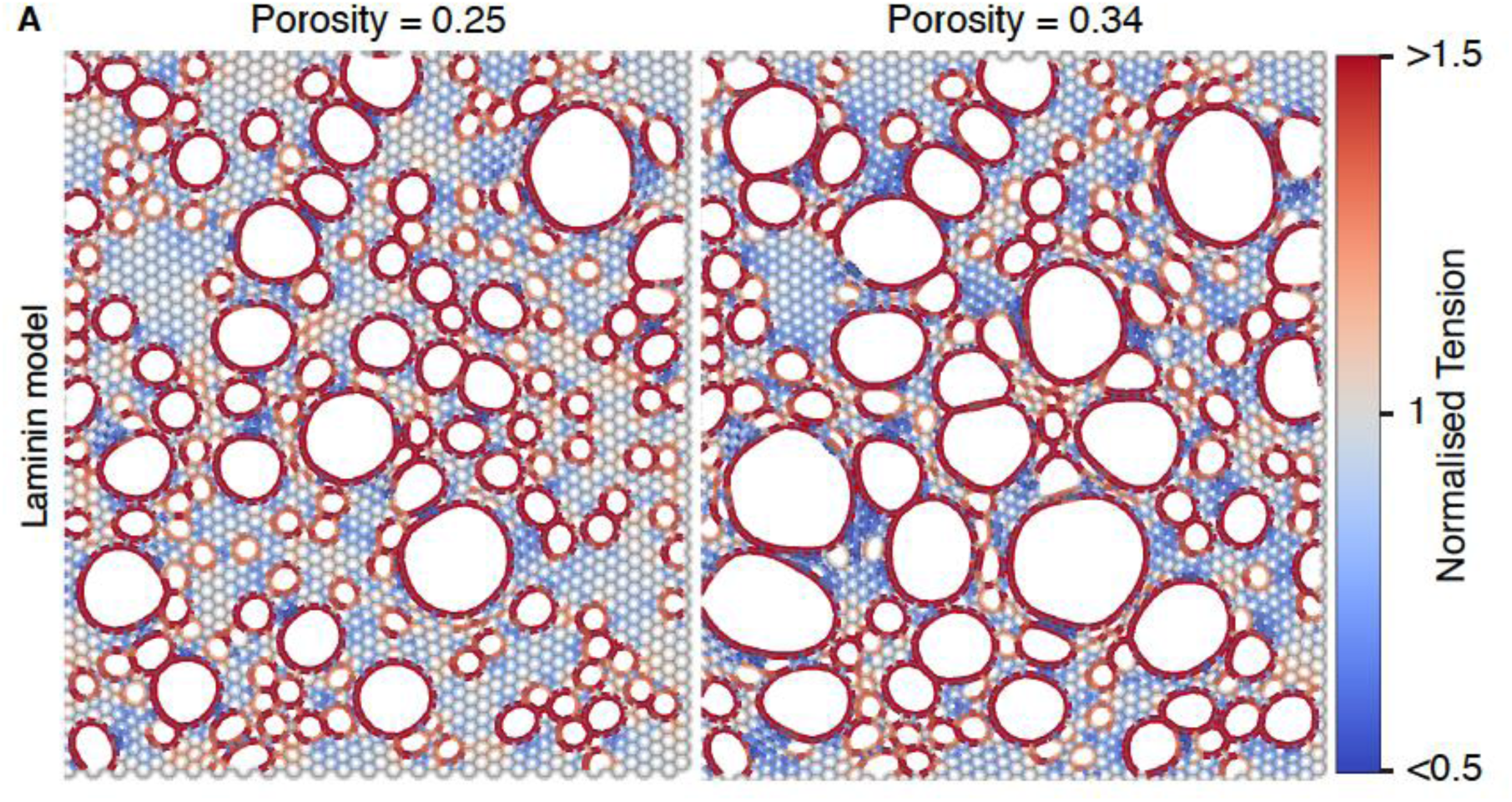
The topology of the lattice is perturbed on greatly increasing the porosity. **(A)** Laminin modeled as a hexagonal lattice with a three-point vertex. Color bar indicates values of normalized tension (tension in porous lattices /tension in a non-porous lattice). Porosity is defined as the ratio of (number of nodes removed/total number of nodes in the network).

**Fig. S5.**
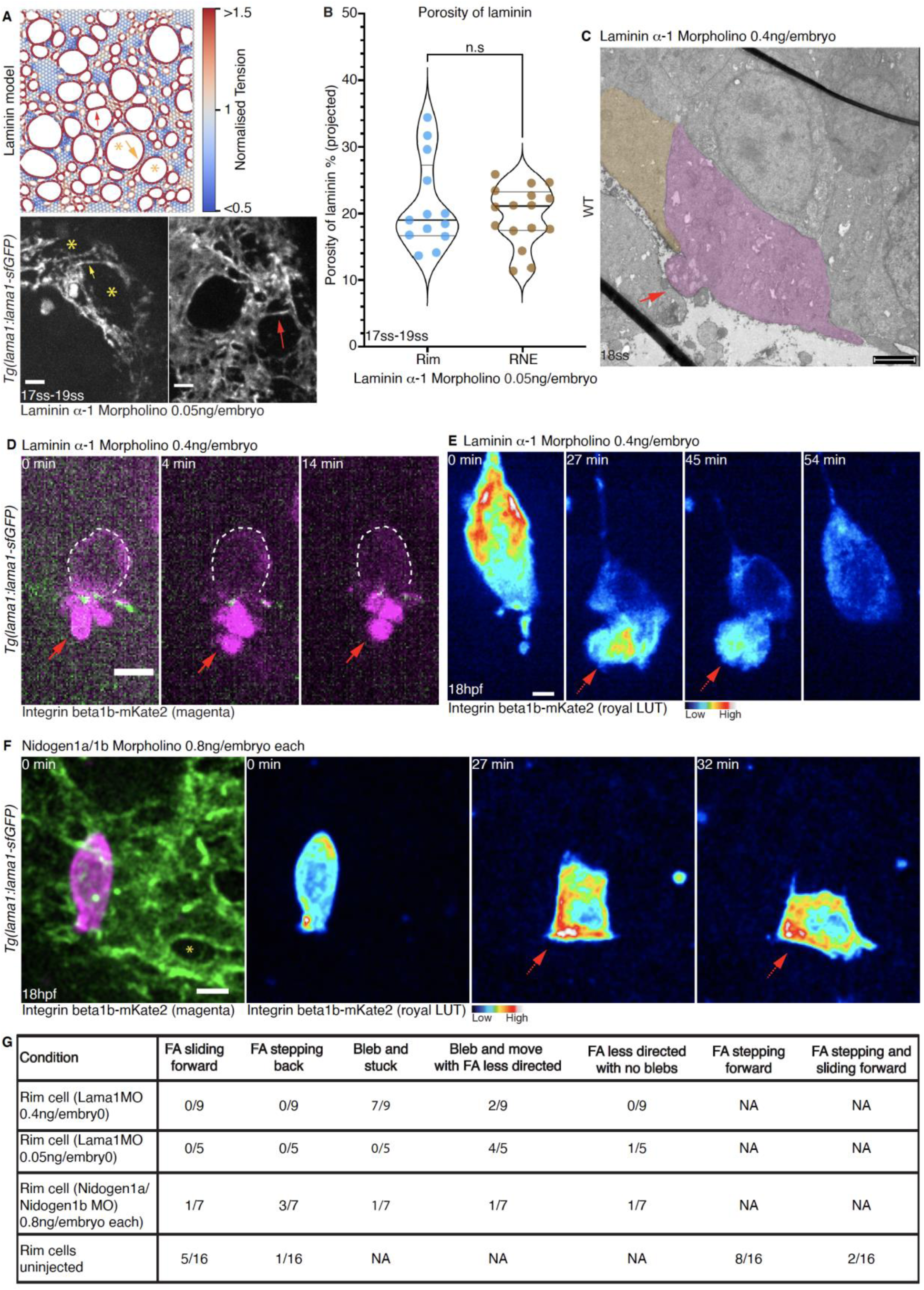
Rim cells exhibit ‘floppy’ behavior upon interference with laminin topology. **(A)** Laminin modeled as a hexagonal lattice with a three-point vertex with a porosity of 0.34. Yellow asterisks, yellow arrows and red arrow point to similarity in structures formed in high porosity lattice models and breaks observed in laminin on treating *Tg(lama1:lama1-sfGFP)* embryos with 0.05ng/embryo laminin α-1 MO (scale bar = 5 µm). Color bar indicates values of normalized tension (tension in porous lattices /tension in a non-porous lattice). Porosity is defined as the ratio of (number of nodes removed/total number of nodes in the network). **(B)** Graph of laminin porosity in the rim (n = 5 embryos) and RNE (n = 7 embryos) region of *Tg(lama1:lama1-sfGFP)* treated with 0.05ng/embryo of laminin α-1 MO. Two tailed Mann Whitney test done (median with quartiles, p = 0.9484). **(C)** TEM image of WT embryo treated with 0.4ng/embryo of laminin α-1 MO. Red arrow points to a bleb (scale bar = 2μm). **(D)** Montage of a rim cell in *Tg(lama1:lama1-sfGFP)* (green) embryo labeled with Integrin beta1b-mKate2 (magenta). Red arrow points to a bleb that is not retracted over 14 minutes (scale bar = 5 μm). **(E)** Montage of a rim cell near the RPE in *Tg(lama1:lama1-sfGFP)* embryos treated with 0.4ng/embryo laminin α- 1 MO and labeled with Integrin beta1b-mKate2 (royal LUT). Dotted red arrow points instances of ‘floppy’ behavior of rim cell in which the rim cell undergoes drastic shape changes (scale bar = 5 μm). **(F)** Montage of rim cell in *Tg(lama1:lama1-sfGFP)* (green) embryos treated with 0.8ng/embryo of Nidogen1a/1b MO each and labeled with Integrin beta1b-mKate2 (in magenta and royal LUT). Yellow asterisk points to the break in laminin. Dotted red arrows point to instances of ‘floppy’ behavior of rim cell (scale bar = 5 μm). **(G)** Table collating the different FA behaviors observed upon perturbing the topology of laminin (NA = ‘non applicable’).

**Fig. S6.**
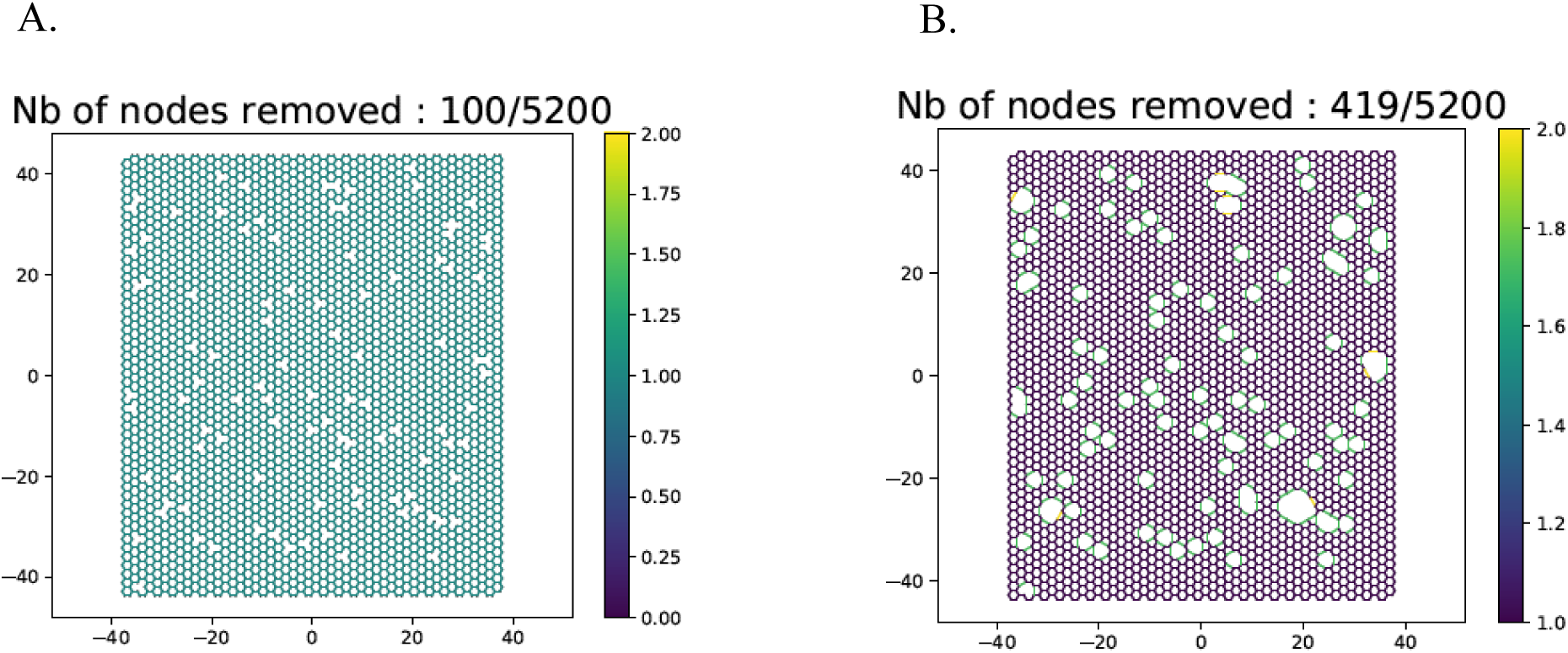
Creating pores in the hexagonal network. **(A)** For n = 100, we randomly remove 100 vertices and their edges. This network consists of vertices with less than 3 neighbors. **(B)** After post-processing, we get a network where all non-boundary vertices have 3 neighbors. Edges of the network are colored by their length.

**Fig. S7.**
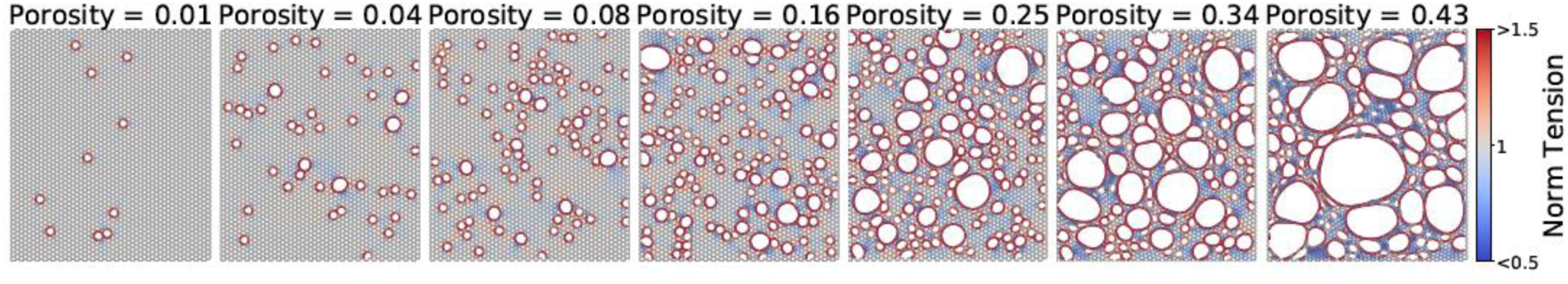
Representative stretched networks of different porosity. Edges are colored by the normalized tension.

**Fig. S8.**
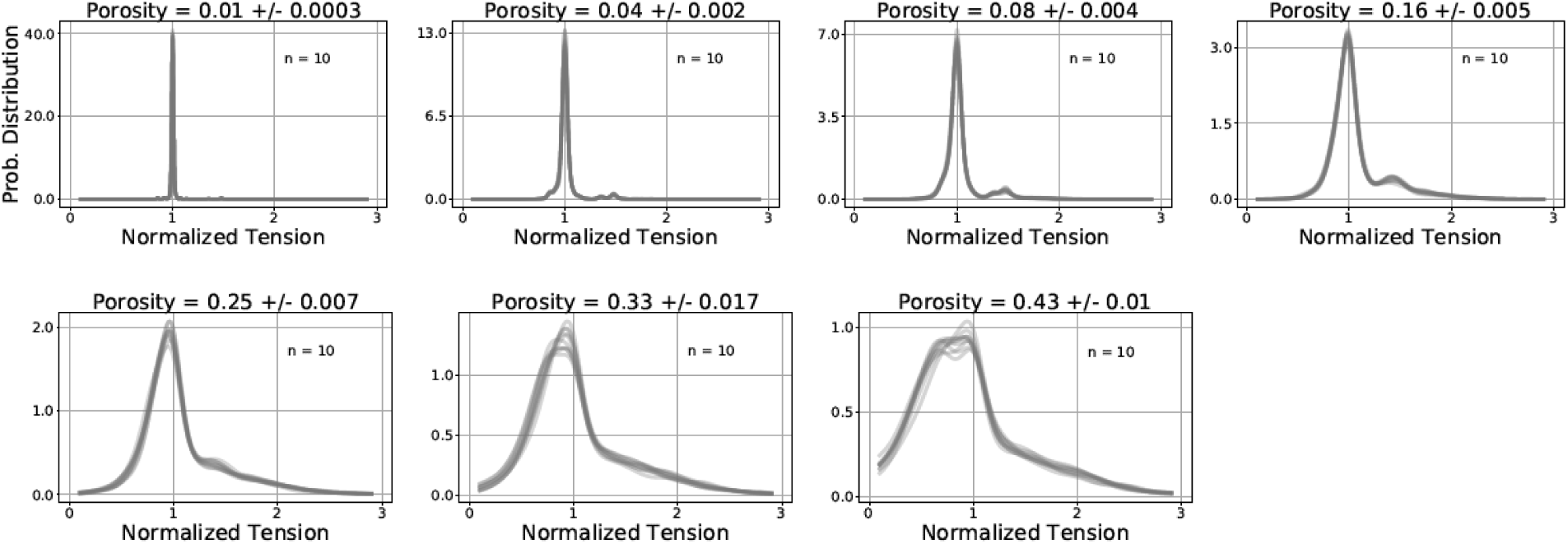
Probability density functions of networks with different porosities. Networks that were generated keeping the same ‘n’ are plotted together where ‘n’ represents the number of nodes removed initially before removing nodes having 2 or fewer number of neighbours.

**Fig. S9.**
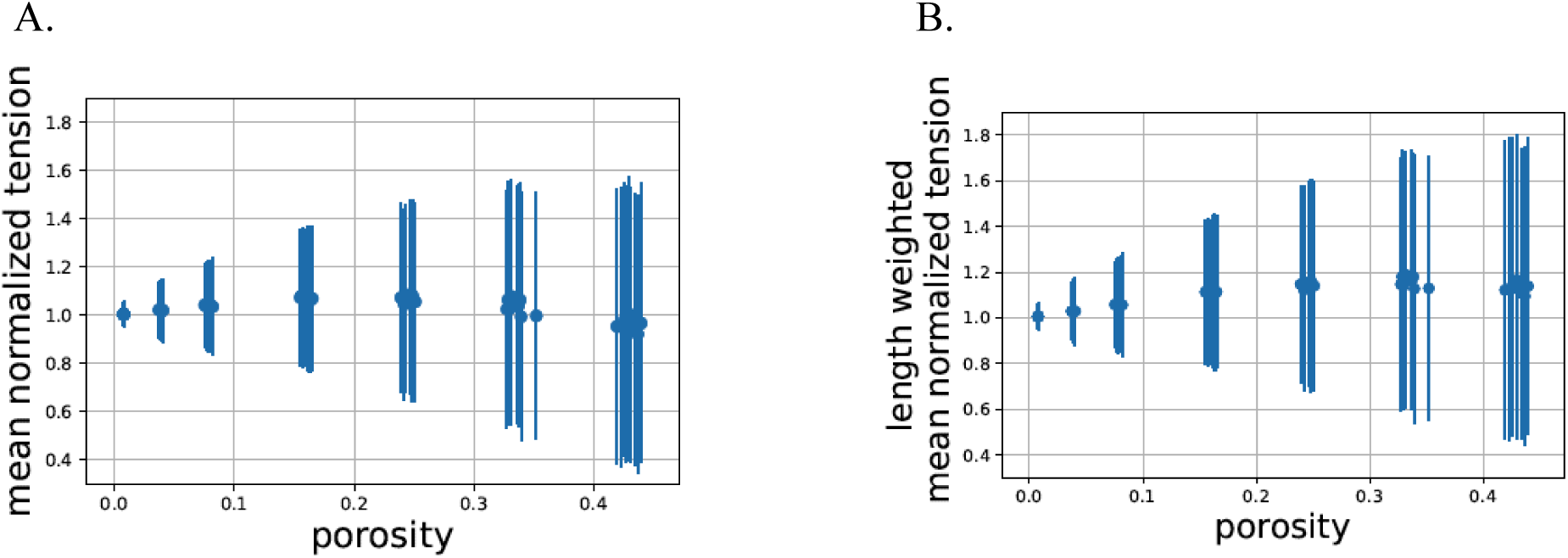
Calculating the normalized tension in networks with differing porosity. **(A)** Normalized tension averaged over all springs in a single network against its porosity is plotted along with the error bars representing the standard deviation in the normalized tension in that network. **(B)** Average of normalized tension weighted by the length of the springs plotted against porosity

**Table S1.**
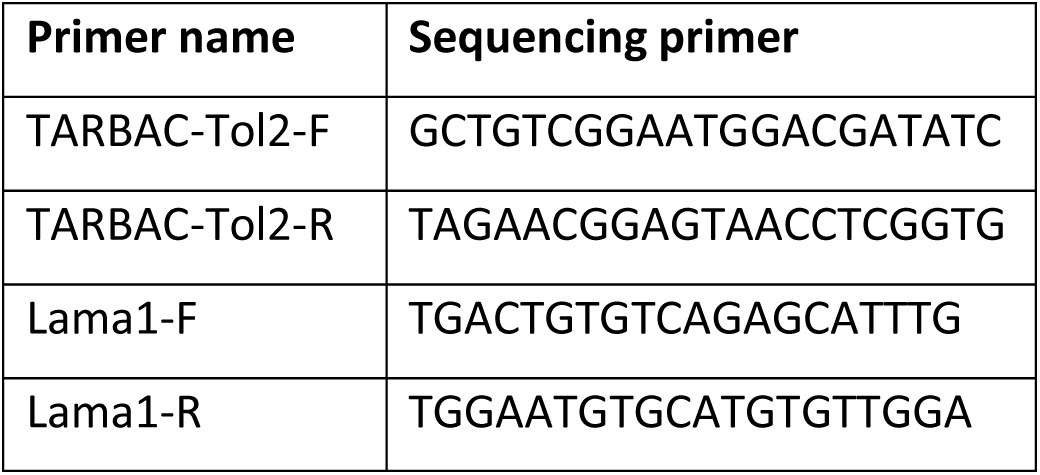
Primers to sequence the lama1 BAC

**Table S2.** Fluorescent protein tags used to label the lama1gene

|  |
| --- |
| 2xTY1-GFP-V5-preTEV-BLRP-FRT-rpsl-Kan-FRT-3xFLAG |
| 2xTY1-mKate2-FRT-rpsl-Kan-FRT-3xFLAG |
| 2xTY1-Dendra-FRT-rpsl-Kan-FRT-3xFLAG |

**Table S3.**
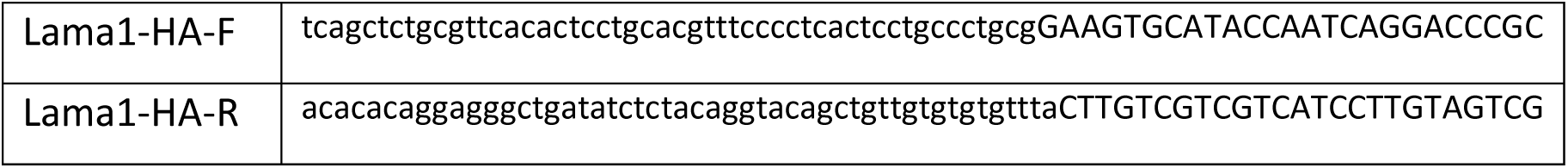
Oligos used for tagging cassettes

**Table S4.** List of antibiotics used at the respective concentrations

| Antibiotic name | Concentration |
| --- | --- |
| Chloramphenicol | 12.5 µg/ml |
| Ampicillin | 100 µg/ml |
| Hygromycin | 100 µg/ml |
| Kanamycin | 15 µg/ml |
| Tetracyclin | 4 µg/ml |

**Table S5.**
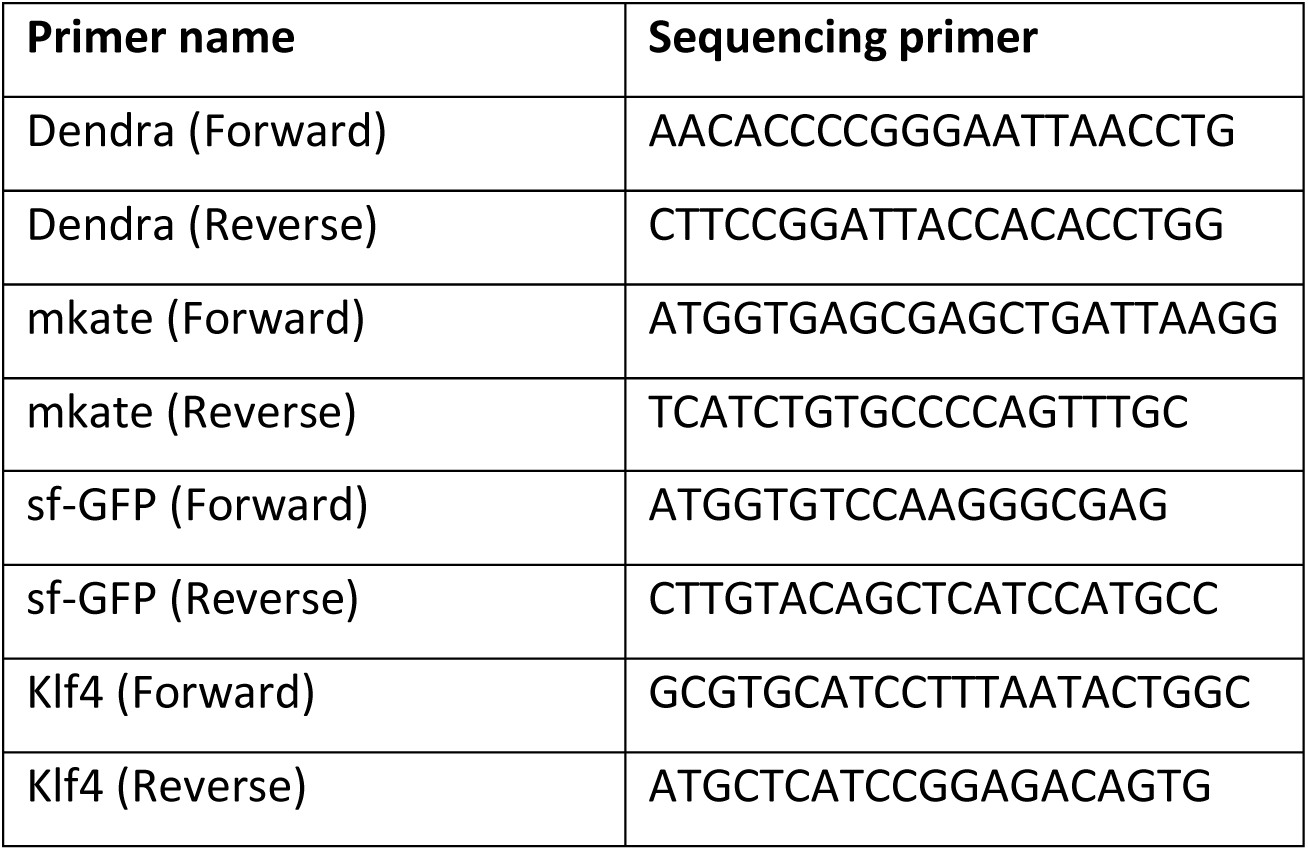
Primers used to genotype the lama1 transgenic lines.

**Table S6.**
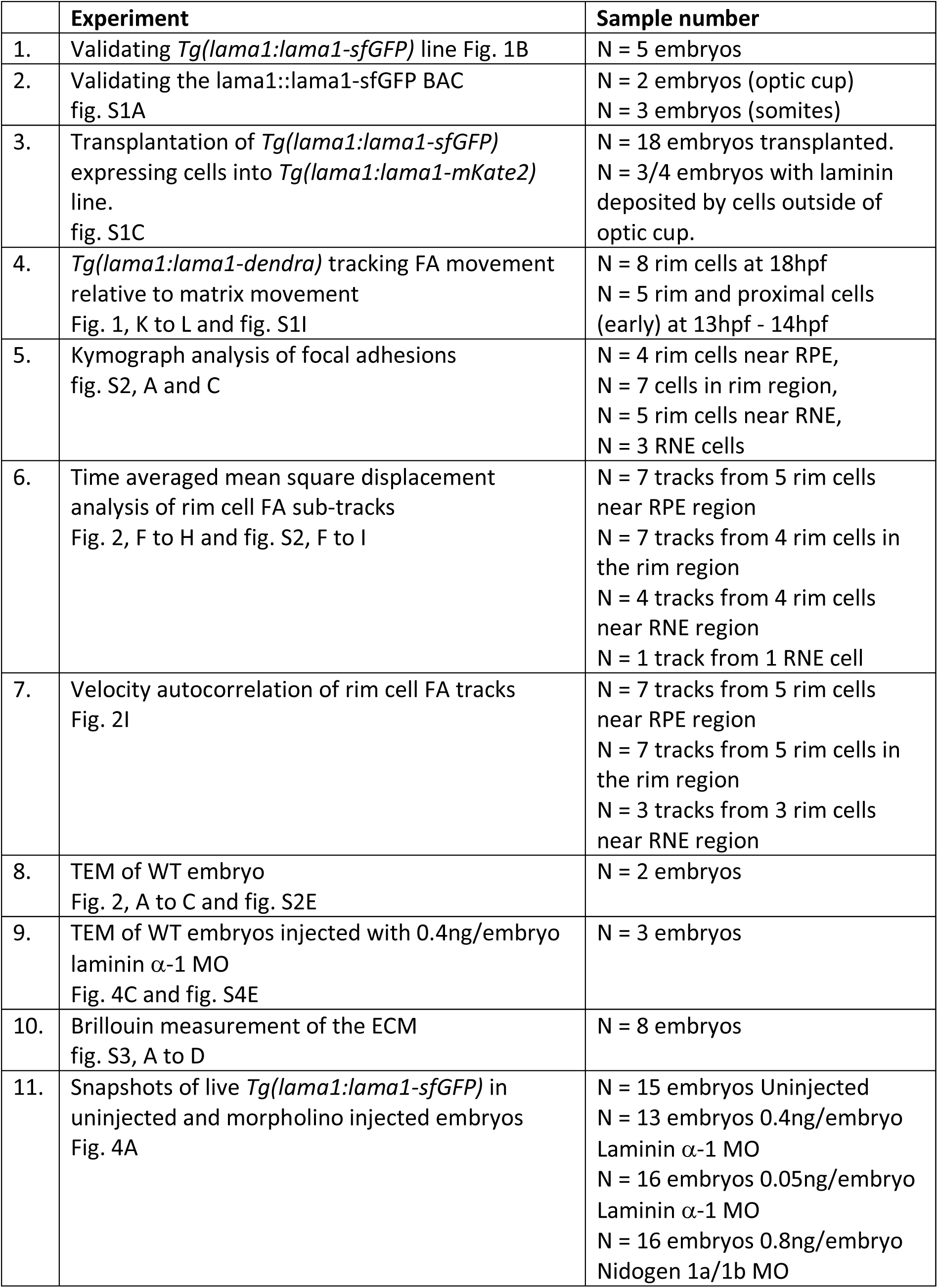
Replicates done for experiments

